# Gut dysbiosis protects against liver injury in autophagy deficient mice by FXR-FGF15 feedback signaling

**DOI:** 10.1101/2020.05.12.090613

**Authors:** Shengmin Yan, Bilon Khambu, Xiaoyun Chen, Zheng Dong, Grace Guo, Xiao-Ming Yin

## Abstract

**Objective:** The gut microbiota (GM) can have complicated and often undetermined interactions with the function of many organs in the body. GM is altered in a variety of liver diseases, but the significance of such changes on the liver disease is still unclear. Hepatic autophagy deficiency causes liver injury accompanied with cholestasis. Here, we investigated the impact of such hepatic changes on GM and in turn the effect of gut dysbiosis on liver injury.

**Design:** Fecal microbiota from mice with liver-specific loss of autophagy-related gene 5 (*Atg5), Atg5*^Δhep^ mice, were analyzed by 16S sequencing. Antibiotics (ABX) was used to modulate GM in mice. Cholestyramine was used to reduce the enterohepatic bile acid (BA) level. The functional role of fibroblast growth factor 15 (FGF15) and ileal farnesoid X receptor (FXR) was examined in mice over-expressing FGF15 gene, or given a fibroblast growth factor receptor 4 (FGFR4) inhibitor.

**Results:** The composition of GM was significantly changed with a notable increase of BA-metabolizing bacteria in *Atg5*^Δhep^ mice, leading to a lower proportion of tauro-conjugated BAs and a higher proportion of unconjugated BAs in the intestine, which markedly activated ileal FXR with an increased expression of FGF15. ABX or cholestyramine treatment exacerbated liver injury and ductular reaction, and decreased FGF15 expression, whereas modulating FGF15 signaling altered liver phenotypes in the autophagy-deficient mice.

**Conclusion:** Gut dysbiosis can remedy liver injury in *Atg5*^Δhep^ mice through the FXR-FGF15 signaling. Antibiotics use in the condition of liver injury may have unexpected adverse consequences via the gut-liver axis.

**SHORT SUMMARY:** *What is already known about this subject?:*

- Gut microbiota (GM) can be altered during hepatic pathogenesis.
- GM are involved in bile acid (BA) metabolism.
- Autophagy deficiency in the liver disrupts BA homeostasis and causes cholestatic injury.

*What are the new findings?:*

- Deficiency of autophagy in the liver causes alteration of GM, which leads to a higher proportion of BA-metabolizing bacteria.
- GM contribute to the activation of ileal farnesoid X receptor (FXR) and a higher expression of fibroblast growth factor 15 (FGF15) in autophagy deficient condition in the liver, which is associated with decreased levels of conjugated BAs and increased levels of unconjugated BAs in the intestine.
- Manipulations that lead to GM alteration, intestinal BA signaling, or FGF15 signaling can all modulate the liver phenotype.
- BA and GM together can act as a sensor to liver injury to trigger FGF15-mediated protective mechanism.

*How might it impact on clinical practice in the foreseeable future?:*

- These findings indicate that gut dysbiosis in the scenario of liver disease can be beneficial, suggesting cautions should be exercised in the use of antibiotics during specific liver diseases.
- If antibiotics need to be used in patients with liver diseases it may be beneficial to enhance the FXR-FGF15 feedback signaling to retain the protective effect of GM.

## INTRODUCTION

Gut microbiota (GM) consist of a diverse community of symbiotic bacteria and have a complex interplay with the host [1]. Interactions between GM and the host result in production of metabolites by microbes, including secondary bile acids (BAs) [1]. Alteration of GM has been associated with multiple diseases, including fatty liver disease. Gut dysbiosis is found in patients with nonalcoholic fatty liver disease (NAFLD) and steatohepatitis (NASH) [2]. In patients with alcohol-related liver disease, the microbiota composition and function are varied in association with the severity of liver condition and whether the patients are active drinkers [3]. On the other hand, gut dysbiosis may contribute to hepatic pathogenesis by translocation of microbial-associated molecular patterns (MAMPs) and live bacteria that can across the gut barrier [4]. However, evidence of beneficial impacts with clear mechanisms is still scarce.

BAs are produced in hepatocytes through two major biosynthetic pathways, including the classic pathway, which converts cholesterol to 7α-hydroxycholesterol (7α-HOC) by cytochrome P450 7A1 (CYP7A1), and the alternative pathway, which converts cholesterol to 27-hydroxycholesterol (27-HOC) by cytochrome P450 27A1 (CYP27A1) [5, 6]. Eventually, the classic pathway leads to the synthesis of cholic acid (CA) and chenodeoxycholic acid (CDCA), whereas the alternative pathway only leads to the synthesis of CDCA in mice [5, 6]. In the human liver, CDCA is an end product, however, CDCA is further converted to muricholic acids (MCAs) by cytochrome P450 2C70 (CYP2C70) in the mouse liver [5, 6]. Primary BAs are conjugated following synthesis and secreted to the intestine, where they are converted by GM into secondary BAs and reabsorbed by the liver through the portal circulation [5, 6]. Dysfunction of BA metabolism and gut dysbiosis can be associated with each other, often in the context of liver diseases [4, 6]. Gut dysbiosis has been found in patients with primary biliary cholangitis or primary sclerosing cholangitis, and in a mouse model of cholestasis [4, 7]. It is not known whether gut dysbiosis would in turn affect liver pathogenesis in the context of dysregulated BA metabolism.

Macroautophagy, hereafter simply referred to as autophagy, is an evolutionarily conserved degradation process and is critical for hepatic homeostasis [8]. Deficiency of key autophagy-related genes (*Atg*) in the liver, e.g. *Atg5* and *Atg7*, causes severe liver injury, fibrosis, and tumorigenesis [9, 10]. Our previous study demonstrates autophagy deficiency-induced liver injury is accompanied with altered BA metabolism, cholestatic injury and impaired hepatic farnesoid X receptor (FXR) activity [11]. We thus hypothesized that autophagy deficiency in the liver could alter GM, which could have further effects on the hepatic phenotype.

Our study shows indeed that hepatic autophagy deficiency leads to an increased proportion of BA-metabolizing bacteria and the alteration of BA composition in the intestine. The significance of these changes is the enhanced activation of ileal FXR and an increased expression of fibroblast growth factor 15 (FGF15). Unexpectedly, we found an exacerbated liver injury and ductular reaction in autophagy-deficient livers following GM removal or blockage of the FGF15-fibroblast growth factor receptor 4 (FGFR4) signaling pathway. Our findings thus indicate that gut dysbiosis can be beneficial, not just being harmful as shown in other studies, in protecting the liver from further injury, and the mechanism is medicated by a gut-liver signaling axis. In addition, the findings imply that antibiotics use in the condition of liver injury should be cautious, considering the potential adverse effect on the liver via the gut-liver signaling.

## METHODS

*Atg5*^F/F^ mice (*B6.129S-Atg5tm1Myok*) [10] and *Atg7*^F/F^ mice [12] had been reported in previous studies. *Atg5*^Δhep^ and *Atg7*^Δhep^ mice were created by crossing *Atg5*^F/F^ or *Atg7*^F/F^ with the Alb:Cre transgenic mice (The Jackson Laboratory, Bar Harbor, ME), respectively. All animal experiments were approved by the Institutional Animal Care and Use Committee (IACUC) of Indiana University. Details of materials, methods, and statistical analysis are included in the Supplementary files.

## RESULTS

### Liver-specific deletion of Atg5 altered the composition of GM

Hepatic autophagy deficiency due to the deletion of a key autophagy gene, *Atg5* or *Atg7*, causes significant liver injury, which is dependent on the nuclear factor erythroid 2-related factor 2 (NRF2) activation [9, 12, 13]. To investigate the impact of hepatic autophagy deficiency on GM, we first assessed the composition of GM in fecal samples from *Atg5*^F/F^ and *Atg5*^Δhep^ mice by 16S sequencing. Principal coordinates analysis showed that the composition of GM was noticeably separated between *Atg5*^F/F^ and *Atg5*^Δhep^ mice at the age of 8- or 16-week (Fig.1A). However, the diversity and the number of species were comparable between *Atg5*^F/F^ and *Atg5*^Δhep^ mice (Fig.1B). The most abundant bacteria at the phylum level were *Bacteroidetes, Firmicutes*, and *Proteobacteria*, and their proportions were in general comparable between *Atg5*^F/F^ and *Atg5*^Δhep^ mice (Fig. 1C). These results suggest that hepatic *Atg5*-deletion alters the proportion of bacterial species rather than the diversity of GM. Analysis at the genus level indeed showed significant disproportions of eight bacteria between *Atg5*^*F/F*^ *and Atg5*^*Δhep*^ *mice* (Fig.1D). A higher proportion of *Lactobacillus* but a lower proportion of *Prevotella, Paraprevotella, Turicibacter, Mogibacterium*, and *Ammonifex* were observed in *Atg5*^Δhep^ mice. The proportion of *Johnsonella* was also increased in most of the *Atg5*^Δhep^ groups except the 16-week old female group. In addition, *Parapedobacter* seemed enriched only in the male *Atg5*^Δhep^ mice but impoverished in the females.

**Fig. 1.**
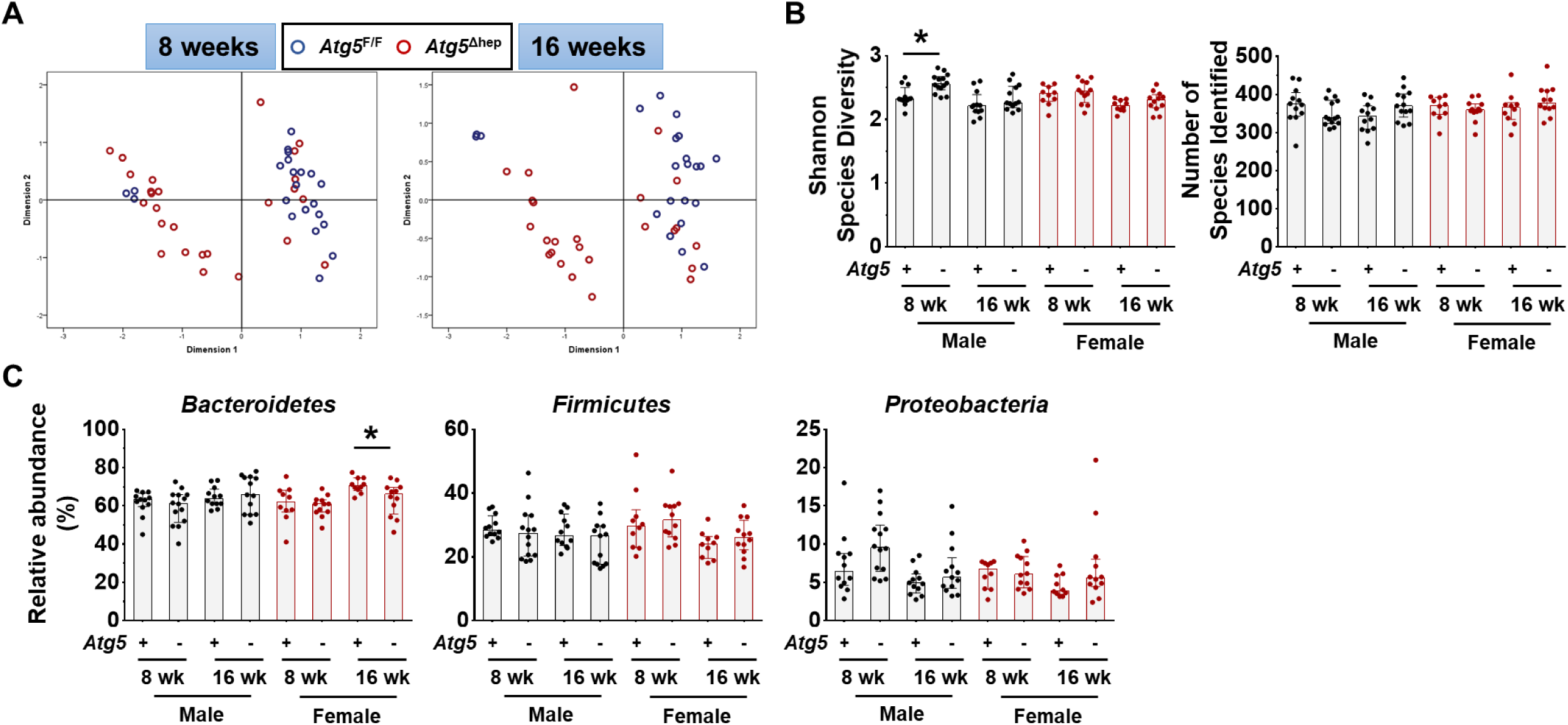
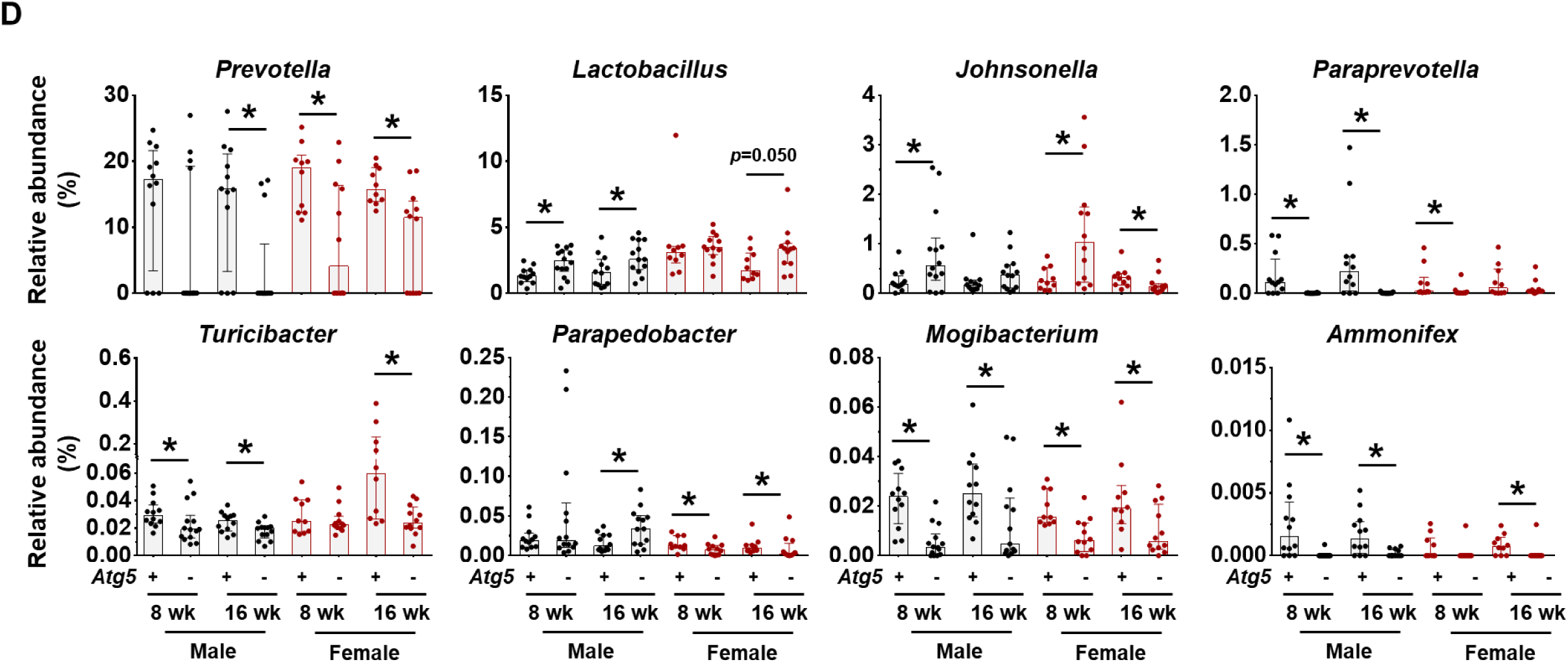
Liver-specific deletion of *Atg5* altered the composition of GM. (**A**). Principal coordinates analysis (PCoA) based on relative abundance at species level shows a distinguishable profile of GM between 8- and 16-weeks old *Atg5*^*F/F*^ and *Atg5*^*Δhep*^ mice. Data from mice of both sexes were plotted. (**B**). Shannon species diversity and number of species identified were similar among different groups of mice. (**C**). Proportion of three most abundant bacteria at phylum level. (**D**). Proportion of bacteria that showed significant changes in at least three comparisons between the age- and sex-matched *Atg5*^*F/F*^ and *Atg5*^*Δhep*^ mice at the genus level. Data were shown as median with interquartile range, n=10/group. Mann-Whitney analysis was used to determine significance, \**p<0.05*.

To understand whether the disproportion of GM was correlated with the autophagy deficiency, we analyzed GM in *Atg7*^Δhep^ mice, another hepatic autophagy-deficiency mouse model. As observed in the *Atg5*^Δhep^ mice, the diversity and the number of identified species were comparable between *Atg7*^Δhep^ mice and their controls (Fig. S1A and B). However, proportions of nine genera were changed in *Atg7*^Δhep^ mice (Fig. S1C), including *Lactobacillus*. These results indicate that hepatic autophagy deficiency can cause gut dysbiosis.

### ABX treatment aggravated Atg5 deficiency-induced liver injury

To investigate the potential impact of GM dysbiosis on the pathogenesis of hepatic autophagy deficiency, mice were given antibiotics (ABX) for 6 weeks (Fig.2A). ABX treatment did not affect either hepatic NRF2 activity (Fig. S2A and B), which could affect liver injury [12], or water consumption (Fig. S2C). The hepatomegaly in *Atg5*^Δhep^ mice remained unchanged following ABX treatment (Fig.2B and C). Nevertheless, gallbladders were enlarged following ABX treatment (Fig.2B and C), consistent with previous findings in germ-free mice [14]. Unexpectedly, serum levels of alanine transaminase (ALT), aspartate transaminase (AST), alkaline phosphatase (ALP), and total BA (TBA) were significantly increased in *Atg5*^Δhep^ mice but not significantly changed in *Atg5*^F/F^ mice following ABX treatment (Fig. 2D). Consistently, ABX treatment enhanced ductular reaction but not fibrotic reaction as shown by Haemotoxylin and Eosin (H&E), cytokeratin 19 (CK19), and Masson’s trichrome staining in *Atg5*-deficient livers (Fig. 2E). Autophagy deficiency causes liver injury accompanied with cholestasis [11]. Consistently, *Atg5*^Δhep^ mice presented increased TBA mainly in the liver and the feces (Fig. 2F). ABX treatment caused further elevations in TBA levels mainly in the intestine and the gallbladder, and thus the total BA pool elevated in *Atg5*^F/F^ mice but more so in *Atg5*^Δhep^ mice (Fig. 2F). Similar to germ-free mice, accumulation of TBA in the intestine was accompanied with decreased fecal levels of TBA following ABX treatment (Fig. 2F). Overall, ABX treatment enhanced cholestatic liver injury of *Atg5*^Δhep^ mice.

**Fig. 2.**
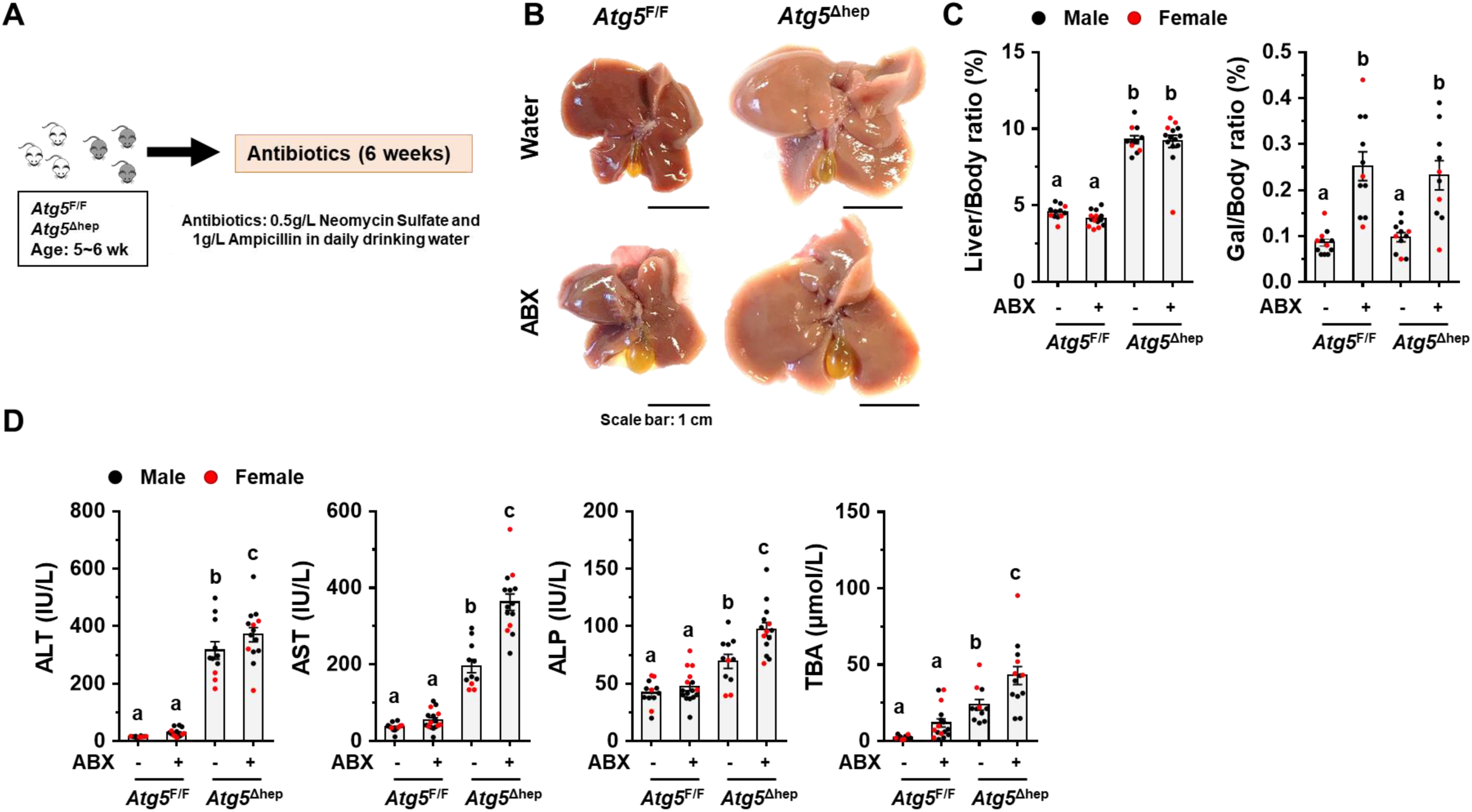
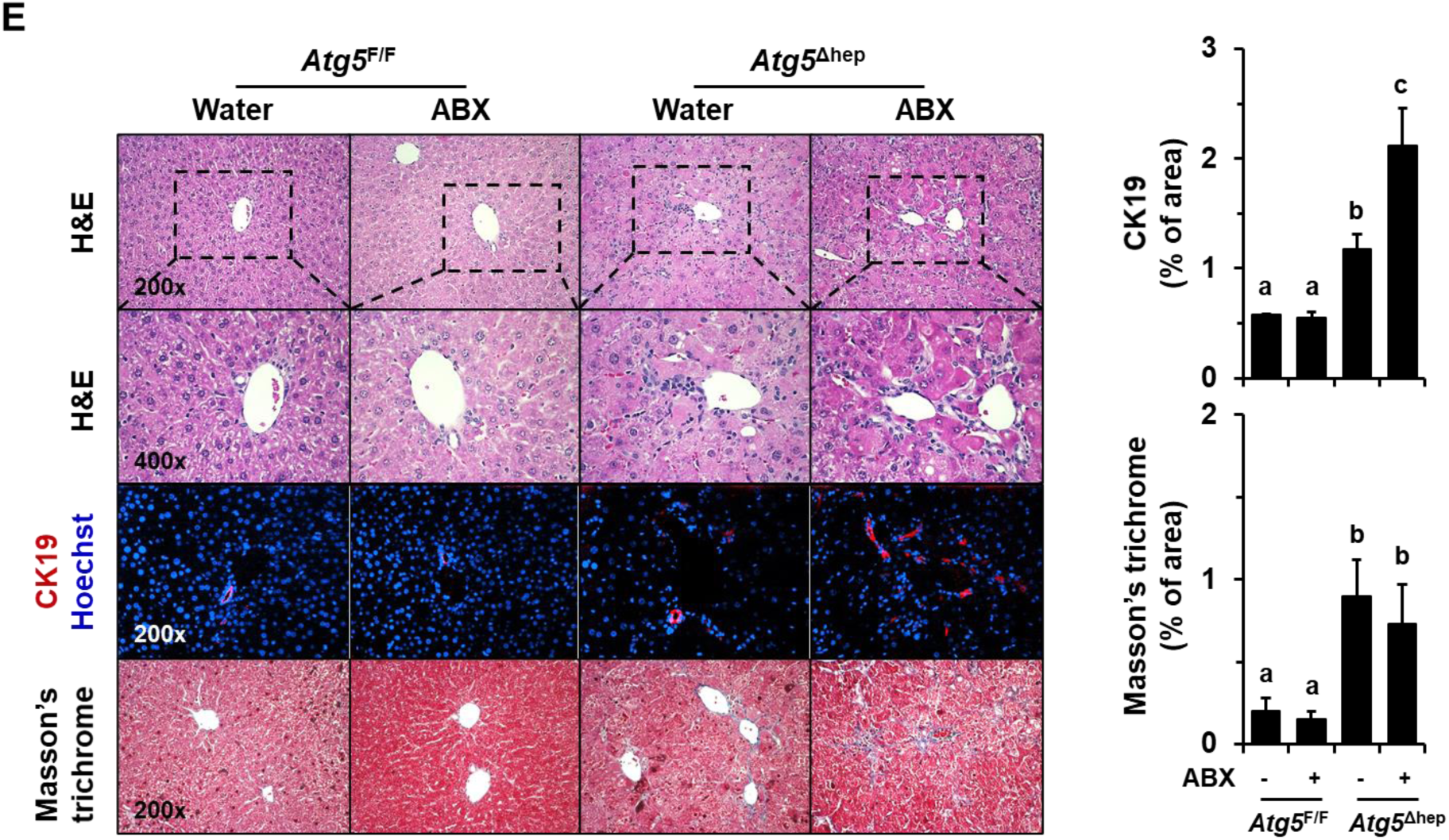
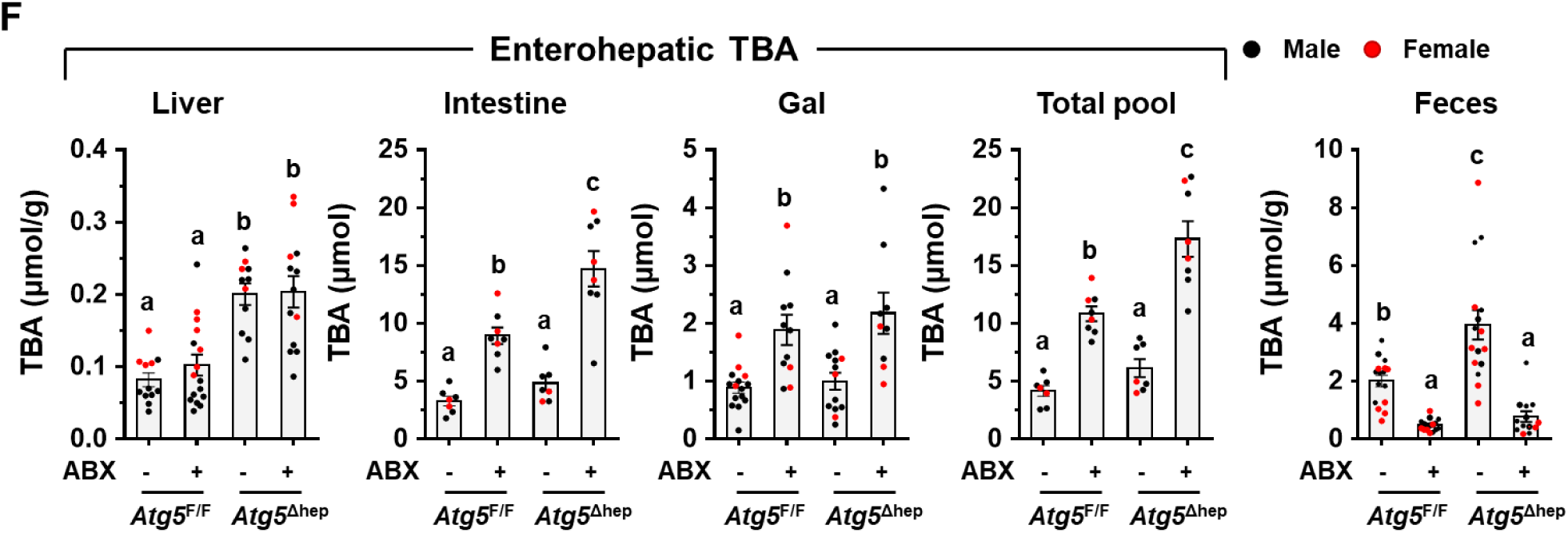
ABX treatment aggravated *Atg5* deficiency-induced liver injury. (**A**). Scheme of the antibiotics (ABX) treatment. Mice were given neomycin sulfate and ampicillin sodium salt mixture in daily drinking water for 6 weeks. (**B**). Representative gross anatomy of livers of indicated genotypes and treatments. (**C**). Liver weight and gallbladder (Gal) weight were determined as percentages of the body weight (n=10-16/group). (**D**). The serum levels of ALT, AST, ALP, and TBA in mice (n=10-16/group). (**E**). Liver sections were subjected to H&E, anti-CK19, or Masson’s trichrome staining. Percentage of positive area was quantified with ImageJ (anti-CK19 staining quantification, n=3-4/group; Masson’s trichrome staining quantification, n=8-12/group). (**F**). TBA levels in indicated compartments were measured (n=7-16/group). Data were shown as means ± S.E. Groups with different letters had significant differences (*p<0.05*). ALT, alanine transaminase; ALP, alkaline phosphatase; AST, aspartate transaminase; CK19, cytokeratin 19; TBA, total bile acids.

In *Atg7*^Δhep^ mice, ABX treatment increased TBA in the intestine and the total pool while decreased fecal excretion of TBA (Fig. S3A), the changes of which are similar to *Atg5*^Δhep^ mice. However, hepatic enzyme levels in the blood were not further elevated in ABX-treated *Atg7*^Δhep^ mice (Fig. S3B). This could be due to the well documented more severe liver injury seen in *Atg7*^Δhep^ mice (Fig. S3B and C v.s. Fig. 2C and D). The more severe phenotype of *Atg7*-deficient livers may mask the effects of ABX treatment.

### *BA-metabolizing bacteria were enriched in Atg5*^Δhep^ *mice*

Since GM are critical for BA metabolism in the intestine [15, 16], we examined whether the disproportion of GM affected BA metabolism in *Atg5*^Δhep^ mice. The major BA-metabolizing bacteria are those that express bile salt hydrolase (BSH) and/or 7α/β-dehydroxylation activity, which include *Lactobacillus, Bacteroides, Bifidobacterium, Eubacterium*, and *Clostridium* [1]. Of note, unlike BSH activity, only a small number of bacteria belonging to the class *Clostridia* have 7α/β-dehydroxylation activity [17]. A higher proportion of *Lactobacillus* (Fig. 1E) but not *Clostridium* or *Eubacterium* (Fig. 3A) was consistently found in *Atg5*^Δhep^ mice. The elevation of *Bacteroides* was only observed in male *Atg5*^Δhep^ mice and that of *Bifidobacterium* was seen only in 16-week old *Atg5*^Δhep^ mice (Fig. 3A). Thus, there were different levels of increments in BA-metabolizing bacteria in *Atg5*^Δhep^ mice. In addition, we also observed an enrichment of *Lactobacillus* at genus level in *Atg7*^Δhep^ mice (Fig. S4A), suggesting that hepatic autophagy deficiency altered proportions of BA-metabolizing bacteria.

**Fig. 3.**
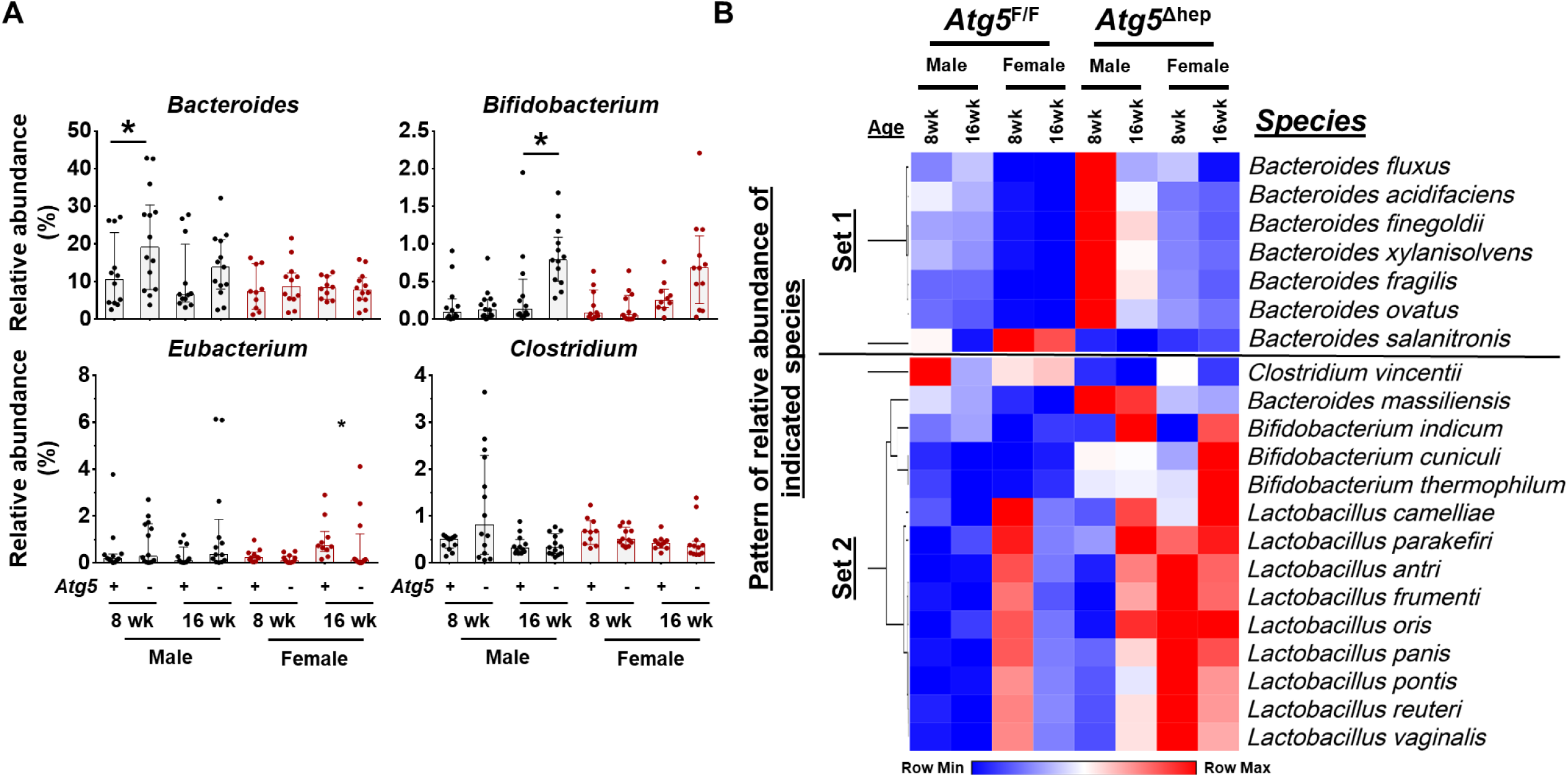
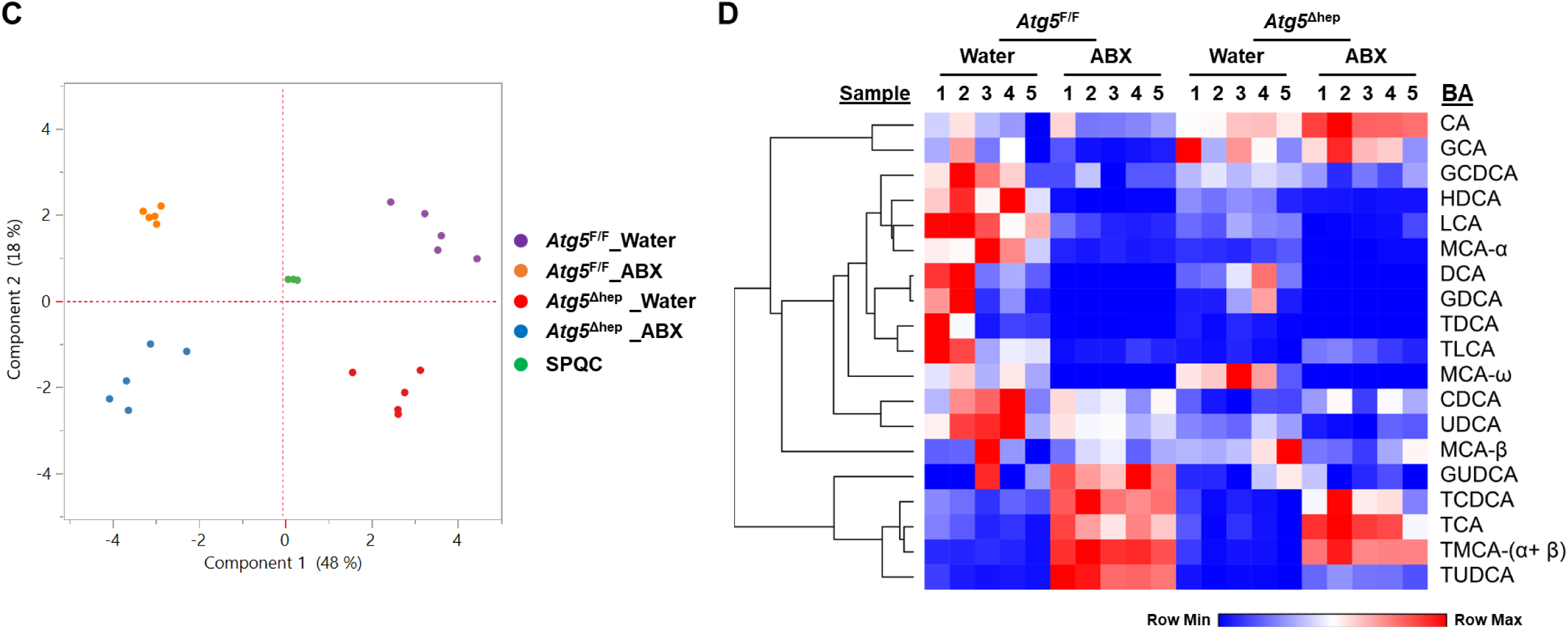
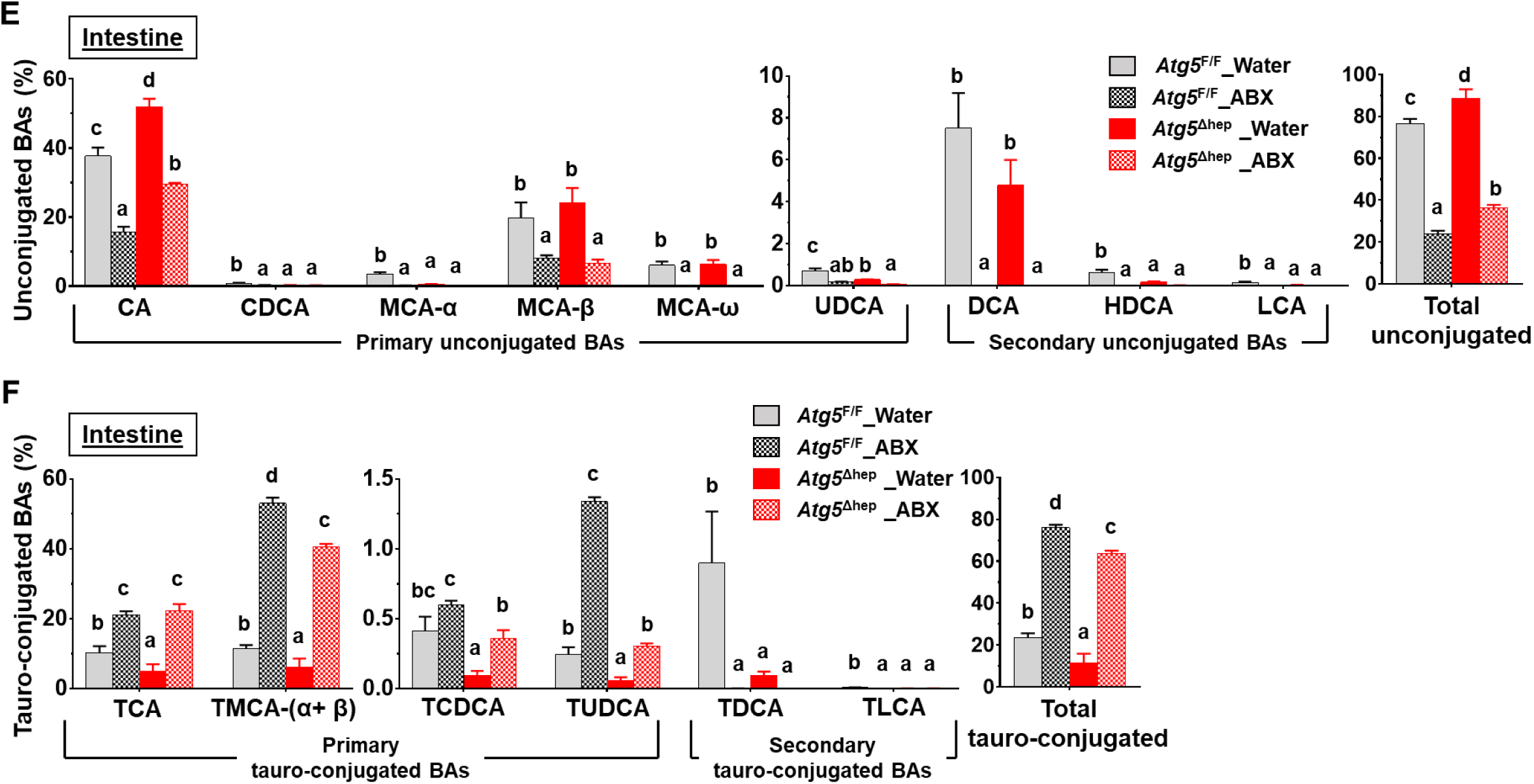
Hepatic autophagy deficiency affect intestinal BA composition in correlation with gut dysbiosis. (**A**). Hepatic autophagy deficiency affected the proportion of bacteria with the bile salt hydrolase and/or 7α/β-dehydroxylation activities at the genus level (*Lactobacillus* is shown in Fig.1E). Data were shown as median with interquartile range, n=10/group. (**B**). The heatmap shows the BA-metabolizing bacteria that are disproportionated in *Atg5*^*Δhep*^ mice at the species level. Heatmap was generated and values in the heatmap were mapped to colors using the minimum and maximum of each row independently. The hierarchical cluster of different species was constructed using one minus Pearson correlation method. Proportion of bacteria in Set 1 was significantly changed in both male and female *Atg5*^*Δhep*^ mice at 8 weeks old. Proportion of bacteria in Set 2 was significantly changed in both male and female *Atg5*^*Δhep*^ mice at 16 weeks old. (**C**). PCoA analysis of BAs in the intestine data (log2-scaled μM). (**D**). Heatmap was generated and values in the heatmap were mapped to colors using the minimum and maximum of each row independently. The heatmap shows the cluster of indicated BA species in the intestine of different groups of mice. The hierarchical cluster of different BAs was constructed using one minus Pearson correlation method. (**E-F**) The intestinal levels of unconjugated (**E**) and tauro-conjugated (**F**) bile acids in male mice. Data were shown as percentage of TBA level (means ± S.E), n=5/group. Groups with different letters or indicated by asterisk had significant differences (*p*<0.05). CA, cholic acid; CDCA, chenodeoxycholic acid; DCA, deoxycholic acid; HDCA, hyodeoxycholic acid; LCA, lithocholic acid; MCA,muricholic acid; TCA, taurocholic acid; TCDCA, taurochenodeoxycholic acid; TDCA, taurodeoxycholic acid; TLCA, taurolithocholic acid; TMCA, tauromuricholic acid; TUDCA, tauroursodeoxycholic acid; UDCA, ursodeoxycholic acid.

Further heterogeneity at the species levels within each of these five genera could be observed in term of the enrichment in *Atg5*^Δhep^ mice. Since the variations could be related to the sex and/or the age, only a few species were overlapped between the cross-age/sex comparisons (Fig. S5A). Two heat maps based on the gender were then generated to include all disproportionated species, which presented the overall disproportion of the BA-metabolizing bacteria in *Atg5*^Δhep^ mice (Fig. S5B). As expected, most of these species with higher proportions were in both male and female *Atg5*^Δhep^ mice, albeit with variations. We then focused on species that were disproportionately altered in both age-matched male and female *Atg5*^Δhep^ mice. We found that among the 7 species that were altered in 8-week old *Atg5*^Δhep^ mice, 6 of them were enriched (Set 1 in Fig. 3B). Similar enrichment of these species was seen in 16-week old *Atg5*^Δhep^ mice although not statistically significant (Fig. 3B). On the other hand, in the 16-week old *Atg5*^Δhep^ mice, 14 other species were disproportionated and 13 of them were enriched (Set 2 in Fig. 3B). We observed similar changes in *Atg7*^Δhep^ mice. At species level, proportion of 9 species, including *Bifidobacterium thermophilum*, and another 8 from genus *Lactobacillus*, was enriched in *Atg7*^Δhep^ mice (Fig. S4B). Overall, our results indicate that BA-metabolizing bacteria are enriched in hepatic autophagy-deficient mice at both genus and species levels, despite that there are age- and sex-related variations.

### Liver-specific deletion of Atg5 altered the composition of intestinal BAs

As BA-metabolizing bacteria were enriched in *Atg5*^Δhep^ mice, we interrogated the composition of BAs in the intestine and the liver. We found that the composition of intestinal BAs was distinctively separated among different groups (Fig. 3C-D). In the intestine, the total level of unconjugated BAs was elevated with a significant increase in CA in *Atg5*^Δhep^ mice (Fig. 3E). Correspondingly, most of the tauro-conjugated BAs, either primary or secondary, were significantly reduced in *Atg5*^Δhep^ mice (Fig. 3F). The composition of intestinal BAs suggested a stronger capacity of deconjugation in *Atg5*^Δhep^ mice. These observations are consistent with the higher proportion of BA-metabolizing bacteria in *Atg5*^Δhep^ mice, which seemed to be more capable of deconjugation, but not necessary dehydroxylation. Following ABX treatment, we observed that the proportion of unconjugated BAs was dramatically increased whereas the proportion of conjugated BAs was decreased in both *Atg5*^F/F^ and *Atg5*^Δhep^ mice (Fig. 3E and F), the pattern of which is similar to the finding in germ-free mice [14]. Interestingly, *Atg5* deletion in the liver did not change the effects of ABX on intestinal composition of BAs, which non-discriminately eliminated BA-metabolizing bacteria for both deconjugation and dihydroxylation capability.

We then examined what potential impact of *Atg5*-deficiency and ABX treatment on hepatic BA compositions as comparisons. BA compositions in the liver were distinctly separable among the groups (Fig. S6A-B). As expected, the major BA species in the liver were the primary conjugated BAs (Fig. S6C-D). Despite total levels of both tauro-conjugated and unconjugated BAs did not seem to be affected significantly, individual BA composition showed changes that were associated with *Atg5* deletion and/or ABX treatment. We observed a significant increase of taurocholic acid (TCA), but not tauromuricholic acid (TMCA), in *Atg5*-deficient livers (Fig. S6D). Hepatic levels of taurochenodeoxycholic acid (TCDCA), tauroursodeoxycholic acid (TUDCA), taurodeoxycholic acid (TDCA), and taurolithocholic acid (TLCA) were all decreased by *Atg5*-deletion (Fig. S6D). Following ABX treatment, hepatic levels of TCA and TCDCA were significantly decreased in *Atg5*^F/F^ but not in *Atg5*^Δhep^ mice (Fig. S6D). Hepatic levels of TMCA-(α+β) and MCA-β were significantly increased in *Atg5*^F/F^ mice, which is similar to the case of germ-free mice [14], whereas the change was not observed in *Atg5*^Δhep^ mice (Fig. S6C-D). These results suggest that hepatic composition of BAs is altered but is less sensitive to ABX treatment in *Atg5*^Δhep^ mice [11].

### Ileal FXR activation and FGF15 expression was upregulated in Atg5^Δhep^ mice in a manner dependent on GM

Intestinal level of TMCA was significantly reduced in *Atg5*^Δhep^ mice, which meanwhile was significantly affected by GM as indicated by the robust elevation following ABX treatment (Fig. 3F). Indeed, TMCA-β in the intestine is known to particularly inhibit ileal FXR, which is activated by TCA [16]. Accumulation of TMCA-β was found in germ-free mice and in conventional mice given ABX treatment, which dramatically inhibited ileal FXR, which thereby reduced expression of FGF15 [14]. FGF15 is secreted into portal circulation to function as a hormone [18]. In the liver, FGF15 can inhibit BA synthesis when binding to its receptor FGFR4, which is critical for BA homeostasis [19, 20, 21, 22, 23].

We therefore hypothesized that the lower level of intestinal TMCA in *Atg5*^*Δhep*^ mice might allow a higher level of ileal FXR activation and a higher level of FGF15 expression. As expected, we found a significant increase of ileal expression of *Fgf15* and small heterodimer partner (*Shp*) in *Atg5*^Δhep^ mice, which is compromised by ABX treatment (Fig. 4A), implying the involvement of TMCA. *Fxr* expression itself was not affected. In addition to *Fgf15* and *Shp*, expression of intestinal BA transporters, which is regulated by FXR [24], is also affected by hepatic autophagy deficiency. In *Atg5*^Δhep^ mice, ileal expression of both ileal bile acid-binding protein (*Ibabp*) and organic solute transporter subunit (*Ost*)*-α*, but not *Ost-β*, was up-regulated, but compromised by ABX treatment (Fig. 4A). Ileal apical sodium–bile acid transporter (*Asbt*) expression was noticeably decreased in *Atg5*^Δhep^ mice (Fig. 4A). Following ABX treatment, expression of *Mrp2* and *Asbt* was induced in mice of both genotypes (Fig. 4A). The pattern of gene expression indicates that the activation of ileal FXR in *Atg5*^Δhep^ mice can be compromised by ABX treatment, suggesting a potential association of these changes with GM.

**Fig. 4.**
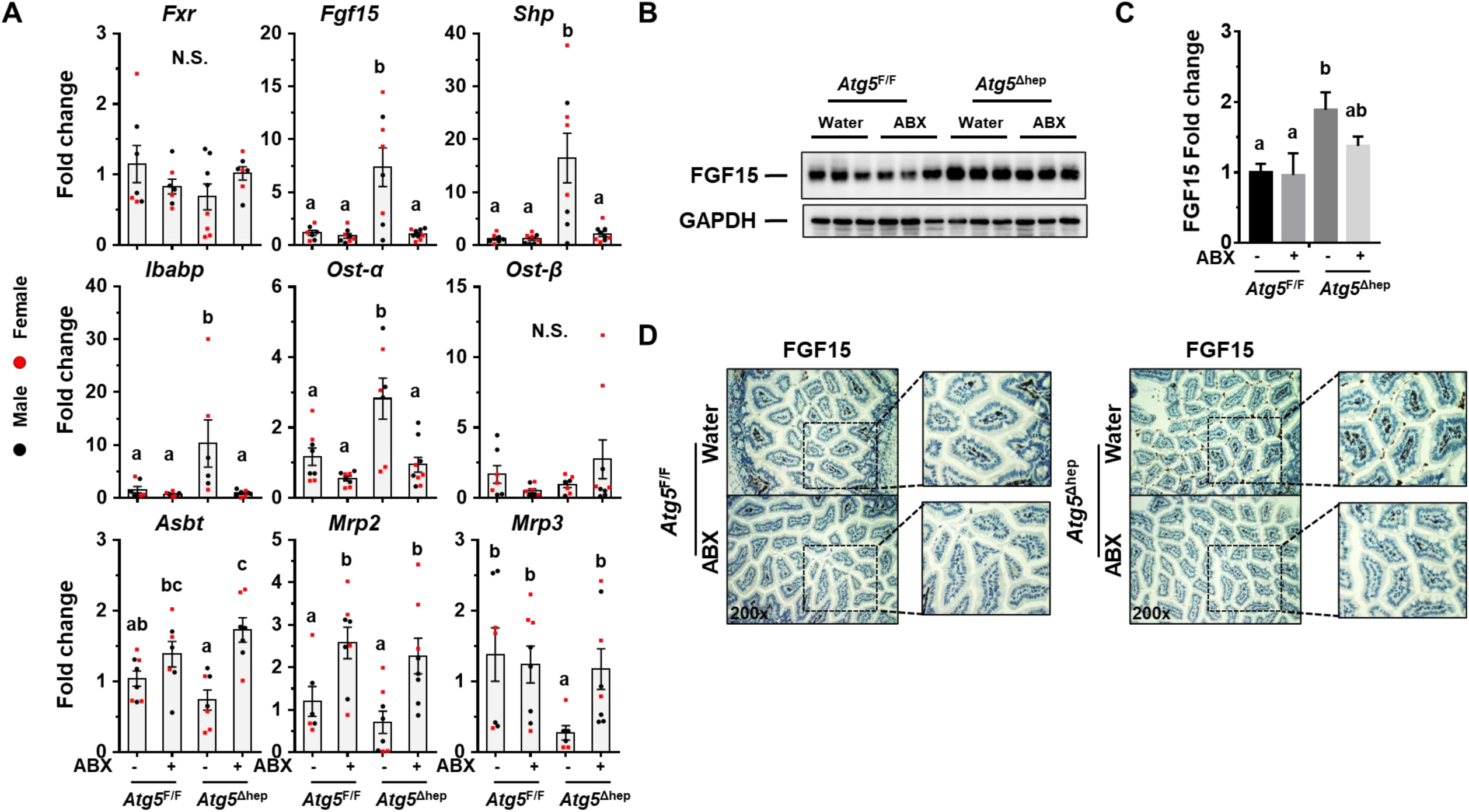
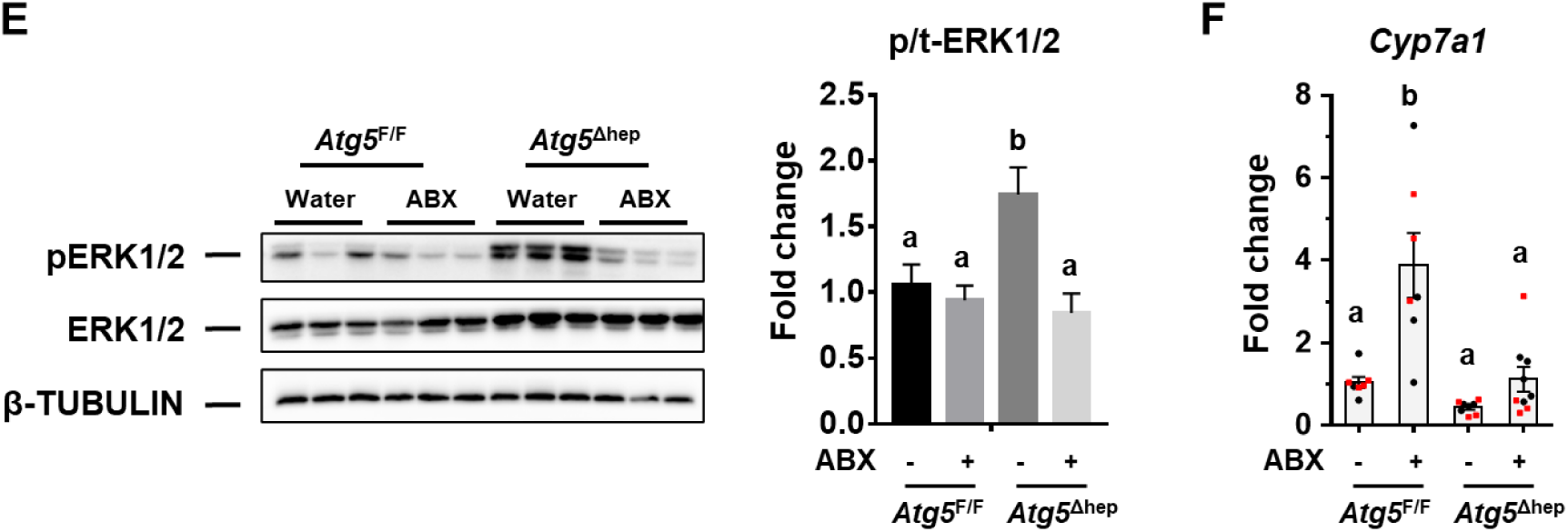
GM regulates ileal FXR activation and FGF15 expression in *Atg5*^*Δhep*^ mice. (**A**). mRNA level of indicated genes in the ileum was analyzed by qRT-PCR (n=7-9/group). (**B-C**). Ileal FGF15 protein levels were examined by immunoblotting assay (**B**), and quantified with densitometry (**C**, n=3/group). (**D**). Representative images of anti-FGF15 immunohistochemistry staining in the ileum of indicated genotypes and treatments. (**E**). ERK1/2 level in the liver was analyzed by immunoblotting assay, and quantified by densitometry. Phosphorylation level of ERK1/2 was normalized by the total protein level and expressed as the fold change of the *Atg5*^F/F^ control group (n=5-7/group). (**F**). Expression of *Cyp7a1* was analyzed by qRT-PCR in the liver (n=6-9/group). Data were shown as means ± S.E. Groups with different letters had significant differences (p<0.05). N.S. indicates no statistical significance. *Asbt* (*Slc10a2*), apical sodium–bile acid transporter; *Cyp7a1*, cytochrome P450 7a1; ERK, extracellular-signal-regulated kinase; *Fgf15*, fibroblast growth factor 15; *Fxr*, farnesoid X receptor; *Ibabp*, ileal bile acid-binding protein; *Mrp2* (*Abcc2*), multidrug resistance-associated protein 2; *Mrp3* (*Abcc3*), multidrug resistance-associated protein 3; *Ost-α* (*Slc51A*), organic solute transporter subunit α; *Ost-β* (*Slc51B*), organic solute transporter subunit β; *Shp*, small heterodimer partner.

The elevation of ileal FGF15 expression was further confirmed by immunoblotting (Fig. 4B and C) and immunochemistry (Fig. 4D), both of which were increased in *Atg5*^Δhep^ mice in a manner dependent on GM. Interestingly, ileal expression of *Fgf15* and *Ibabp* was induced in *Atg7*^Δhep^ mice, which was also compromised following ABX treatment (Fig. S7), suggesting that the modulation of ileal FXR activity by hepatic autophagy deficiency was not dependent on specific autophagy-related genes.

In order to examine the consequent impacts of FGF15 on *Atg5*-deficient livers, we analyzed FGF15-related pathways. In livers, FGF15 can bind to FGFR4 and activate the extracellular-signal-regulated kinase (ERK) pathway [19]. As expected, phosphorylation level of ERK1/2 in *Atg5*-deficient livers was significantly induced but compromised following ABX treatment (Fig. 4E). *Cyp7a1* gene that encodes the rate-limiting enzyme in the classical BA synthesis pathway is negatively regulated by FGF15 signaling [19]. Indeed, while *Cyp7a1* mRNA level is low in *Atg5*^Δhep^ livers, its expression was noticeably elevated following ABX treatment (Fig. 4F), which is similar to the case in the germ-free mice [14] and consistent with the alteration in the FGF15 signaling. Expression of *Fxr* and most of its target genes was suppressed in *Atg5*-deficient liver as shown in our previous study [11] but not affected by ABX treatment (Fig. S8A-B).

Taken together, ABX treatment affects the intestinal composition of BA, and eliminates the relative advantage of *Atg5*^Δhep^ mice in producing more ileal FGF15 due to a lower level of TMCA in the intestine.

### BAS reduced TBA pool but also ileal FGF15 production, leading to an enhanced Atg5 deficiency-induced liver injury

Bile acid sequestrants (BAS) are large polymers that bind negatively charged bile acids in the small intestine, which can prevent reabsorption of bile acids in the gut, increase their fecal excretion, and finally disrupt enterohepatic circulation [25]. BAS is efficient to reduce TBA pool, however, evidence also shown that it can consequently inhibit ileal FXR and reduce expression of FGF15 in mice [26] and FGF19 in human [19]. To determine the impact of enterohepatic TBA on autophagy-deficient livers, mice were given cholestyramine resin treatment (Fig. 5A). As expected, a significant decrease of TBA levels in livers and enterohepatic circulation (Fig. 5B), and reduced gallbladder size (Fig. 5C) were observed in mice following BAS treatment. Although hepatomegaly in *Atg5*^Δhep^ mice was not further enhanced (Fig. 5C), serum levels of ALT, AST, and ALP were all increased in *Atg5*^Δhep^ mice following BAS treatment (Fig. 5D). H&E staining showed an increased number of oval cells around the periportal areas in *Atg5*-deficient livers following BAS treatment (Fig. 5E). The increased positive areas of CK19 and Masson’s trichrome staining indicate a more severe ductular reaction in *Atg5*-deficient livers following BAS treatment (Fig. 5E). These results suggest that BAS treatment in *Atg5*^*Δhep*^ mice paradoxically enhances liver injury.

**Fig. 5.**
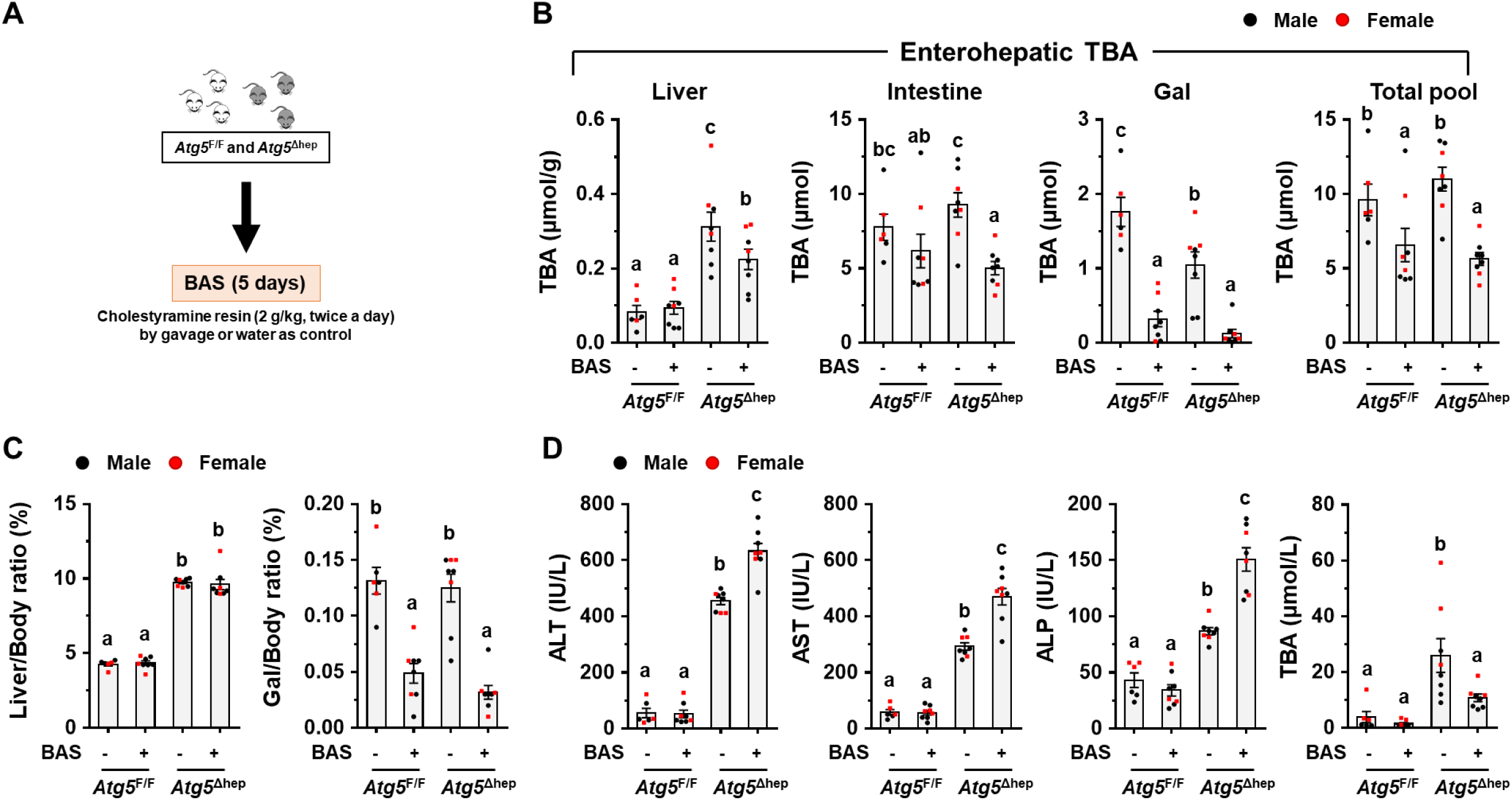
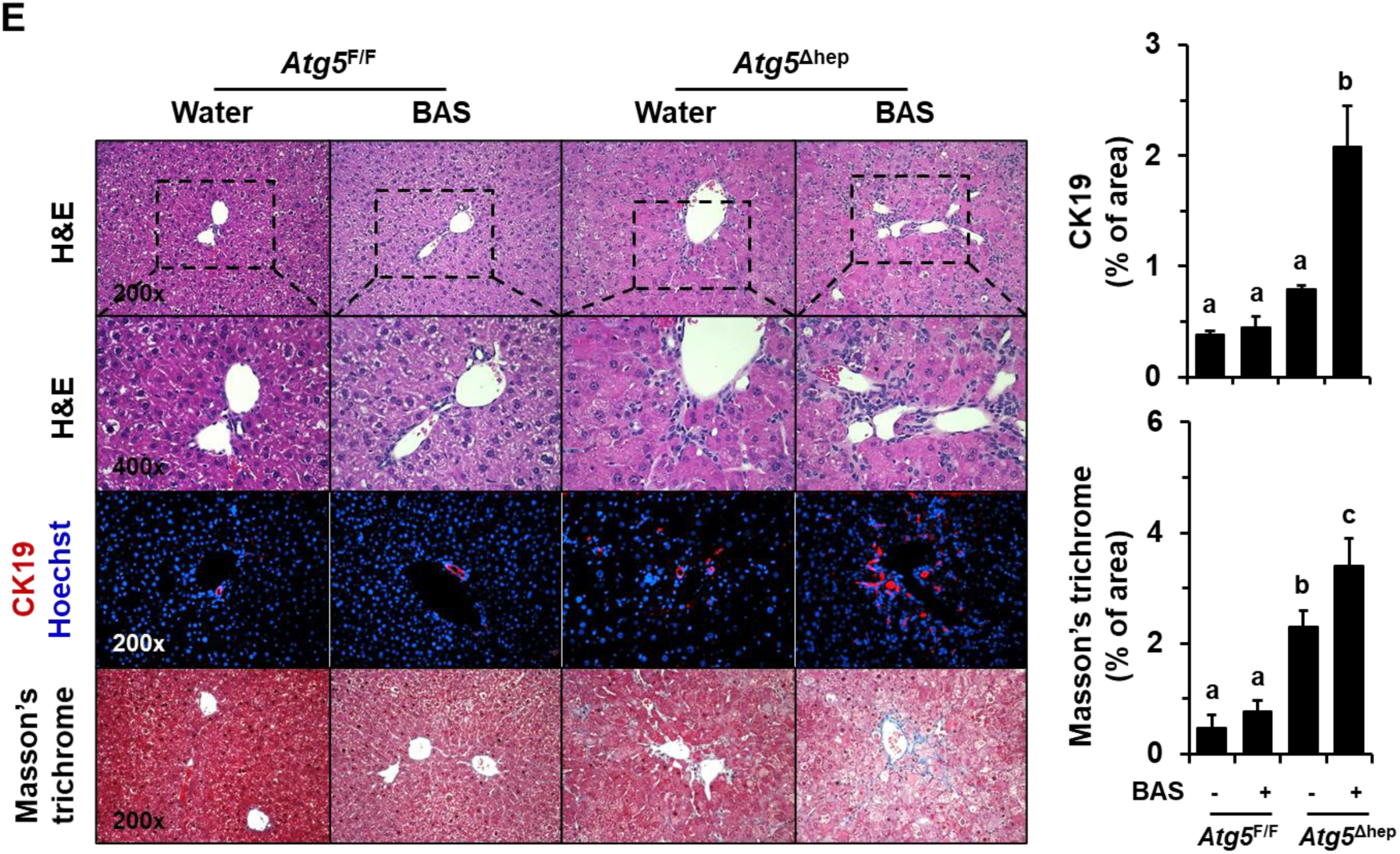
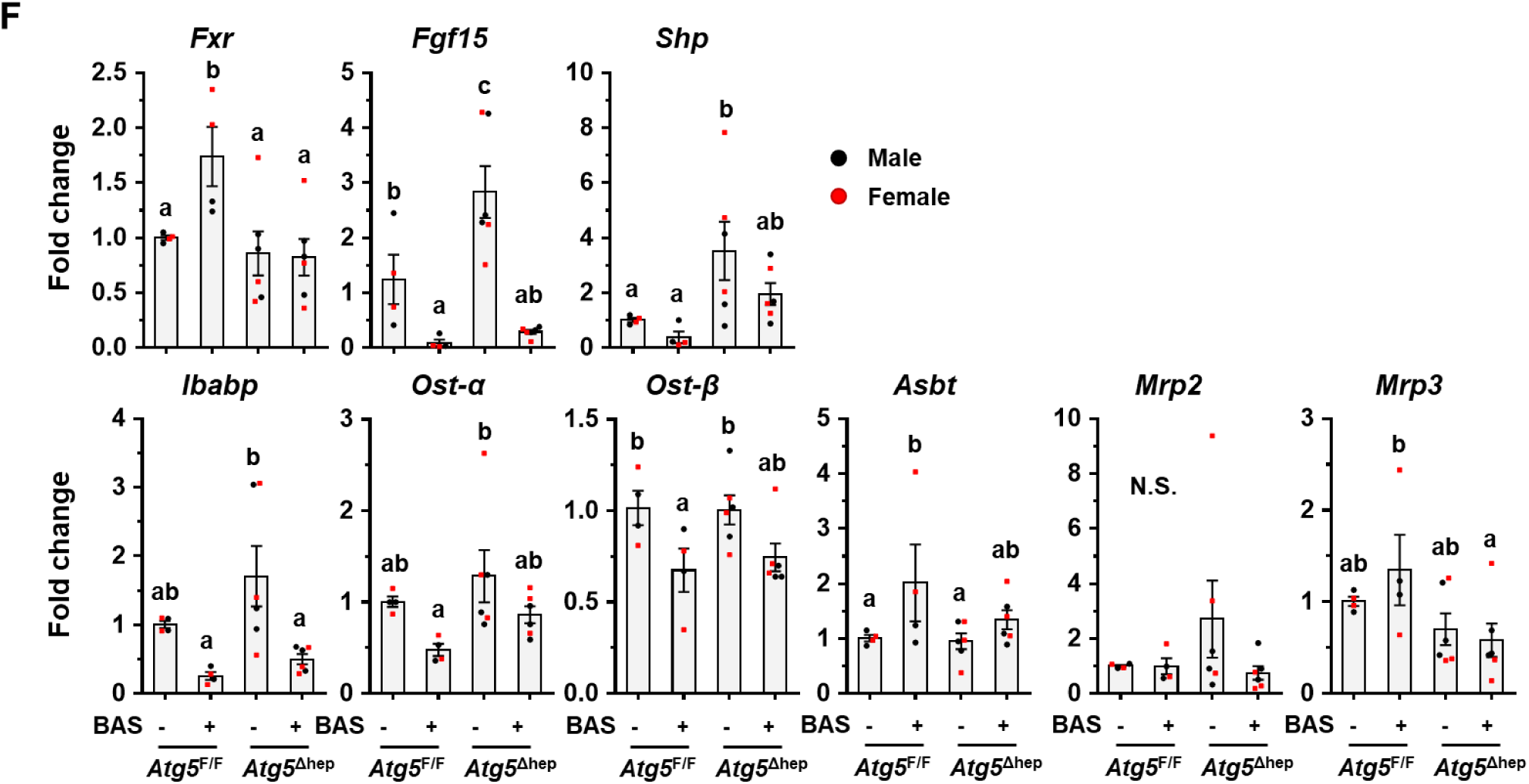
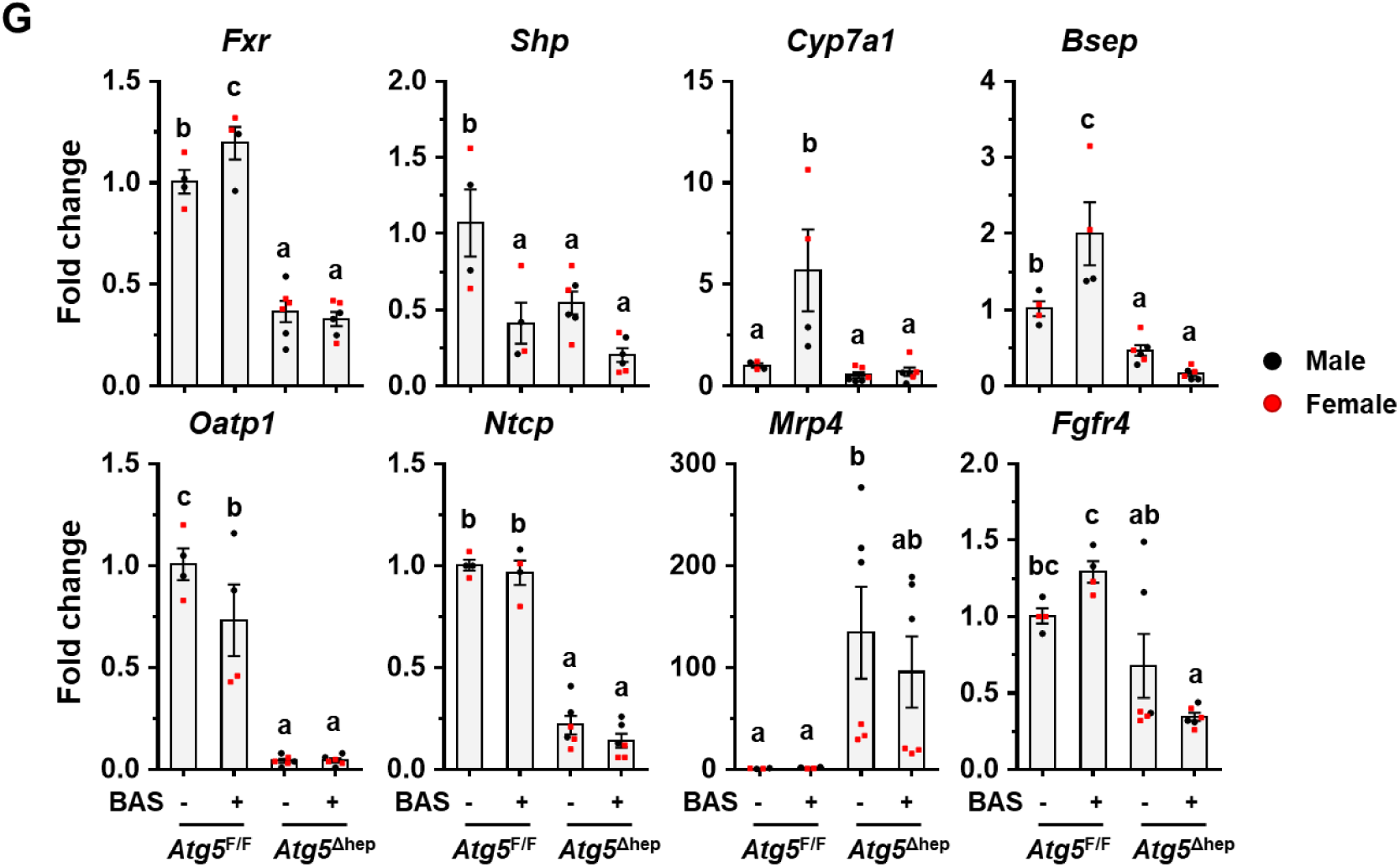
BAS reduced TBA pool but enhanced *Atg5* deficiency-induced liver injury. (**A**). Scheme of the bile acid sequestrant (BAS) treatment. Water was given as control. (**B**). TBA levels in the indicated compartments were measured (n=6-8/group). (**C**). Liver weight and gallbladder (Gal) weight were determined as percentage of the body weight (n=6-8/group). (**D**). The serum levels of ALT, AST, ALP, and TBA in mice following BAS treatment (n=6-8/group). (**E**). Liver sections were subjected to H&E, anti-CK19, or Masson’s trichrome staining. Percentage of positive area was quantified with ImageJ (anti-CK19 staining quantification, n=3-4/group; Masson’s trichrome staining quantification, n=3-5/group). (**F-G**). Expression of genes related to BA metabolism in ileums (**F**) or in livers (**G**) was analyzed by qRT-PCR (n=4-6/group). Data were shown as means ± S.E. Groups with different letters had significant differences (*p<0.05*). ALT, alanine transaminase; ALP, alkaline phosphatase; AST, aspartate transaminase; *Asbt* (*Slc10a2*), apical sodium–bile acid transporter; *Bsep* (*Abcb11*), bile salt export pump; *Cyp7a1*, cytochrome P450 7a1; CK19, cytokeratin 19; *Fgf15*, fibroblast growth factor 15; *Fgfr4*, fibroblast growth factor receptor 4; *Fxr*, farnesoid X receptor; *Ibabp*, ileal bile acid-binding protein; *Mrp2* (*Abcc2*), multidrug resistance-associated protein 2; *Mrp3* (*Abcc3*), multidrug resistance-associated protein 3; *Mrp4* (*Abcc4*), multidrug resistance-associated protein 4; *Ntcp* (*Slc10a1*), Na/Taurocholate cotransporting polypeptide; *Oatp1* (*Slco1a1*), organic anion-transporting polypeptide 1; *Ost-α* (*Slc51A*), organic solute transporter subunit α; *Ost-β* (*Slc51B*), organic solute transporter subunit β; *Shp*, small heterodimer partner; TBA, total bile acids.

In order to identify potential mechanisms in which BAS treatment contributed to enhanced liver injury in *Atg5*^*Δhep*^ mice, we examined expression of FXR-regulated genes in both ileums and livers. Despite an increase of *Fxr* gene expression in *Atg5*^F/F^ mice following BAS treatment, ileal expression of FXR-promoted genes, including *Fgf15, Shp, Ibabp, Ost-α*, and *Ost-β*, was significantly reduced in mice following BAS treatment (Fig. 5F). The ileal level of *Asbt* mRNA, an FXR-suppressed gene, was increased following BAS treatment (Fig. 5F). These results suggested that ileal FXR activity was strongly inhibited by BAS treatment and consequently decreased expression of ileal *Fgf15*. In the liver, BAS treatment suppressed hepatic FXR activity in *Atg5*^F/F^ mice but did not significantly change the expression of FXR targets in *Atg5*^*Δhep*^ mice (Fig. 5G), perhaps because the FXR activity was already reduced in autophagy deficient livers. Taken together, these results suggest that BAS treatment in *Atg5*^*Δhep*^ mice reduces ileal FXR activation and *Fgf15* expression, which may contribute to the paradoxical effects on autophagy deficiency-induced liver injury.

### Intestine-specific FXR agonist activated ileal FXR in Atg5^F/F^ mice but not in Atg5^Δhep^ mice

Since both ABX and BAS treatments exacerbate liver injury in *Atg5*^Δhep^ mice accompanied with reduced ileal FXR activity, we asked whether further activate ileal FXR can improve *Atg5* deficiency-induced liver injury. Hence, mice were given an intestine-specific FXR agonist, Fexaramine (FEX), for 7 days (Fig. S9A). Unexpectedly, FEX treatment did not change serum levels of ALT, AST, ALP, and TBA in *Atg5*^Δhep^ mice, whereas serum level of TBA was decreased in *Atg5*^F/F^ mice (Fig. S9B). Analysis of the expression of ileal *Fxr* and its target genes suggested an activation of FXR in *Atg5*^F/F^ mice but not in *Atg5*^Δhep^ mice following FEX treatment (Fig. S9C). The TBA pool and fecal TBA level were both significantly reduced in *Atg5*^F/F^ mice following FEX treatment, but not so much in *Atg5*^Δhep^ mice (Fig. S9D). The results suggest that altered GM and saturated FXR with existing BA agonists in *Atg5*^Δhep^ mice may reduce the effects of FEX treatment in these mice. An early study also showed that GM could contribute to the efficacy of FEX treatment [27].

### Overexpression of FGF15 attenuated pathological features in Atg5^Δhep^ livers

FGF15 was significantly induced in *Atg5*^Δhep^ mice but was reduced following treatments with ABX or BAS. We thus hypothesized that an elevated level of FGF15 could play a protective role in *Atg5* deficiency-induced liver injury. Since recombinant FGF15 does not seem stable in the circulation for long-term experimentation [28, 29], AAV-FGF15 was given to *Atg5*^Δhep^ mice (Fig. 6A). The level of phosphorylated ERK1/2 in the liver was elevated following AAV-FGF15 injection 1 week and 4-weeks later (Fig. 6B), suggesting that FGF15 signaling was enhanced in the liver as supported by a higher hepatic expression of *Fgf15* (Fig. 6C). Consequently, hepatic levels of two classic targets of FGF15 signaling, *Cyp7a1* and cytochrome p450 family 8b1 (*Cyp8b1*), were reduced following FGF15 overexpression (Fig. 6C). Paradoxically, hepatic *Fxr* and *Shp* expression was not elevated following FGF15 overexpression although the expression of bile salt export pump (*Bsep*), a direct target of FXR, was induced (Fig.6C), suggesting FXR-independent effects of FGF15 on hepatic gene expression. Consistent with the reduced expression of *Cyp7a1* and *Cyp8b1*, the total TBA pool, mainly contributed by the intestinal level of BA, and the fecal TBA level were significantly reduced (Fig. 6D).

**Fig. 6.**
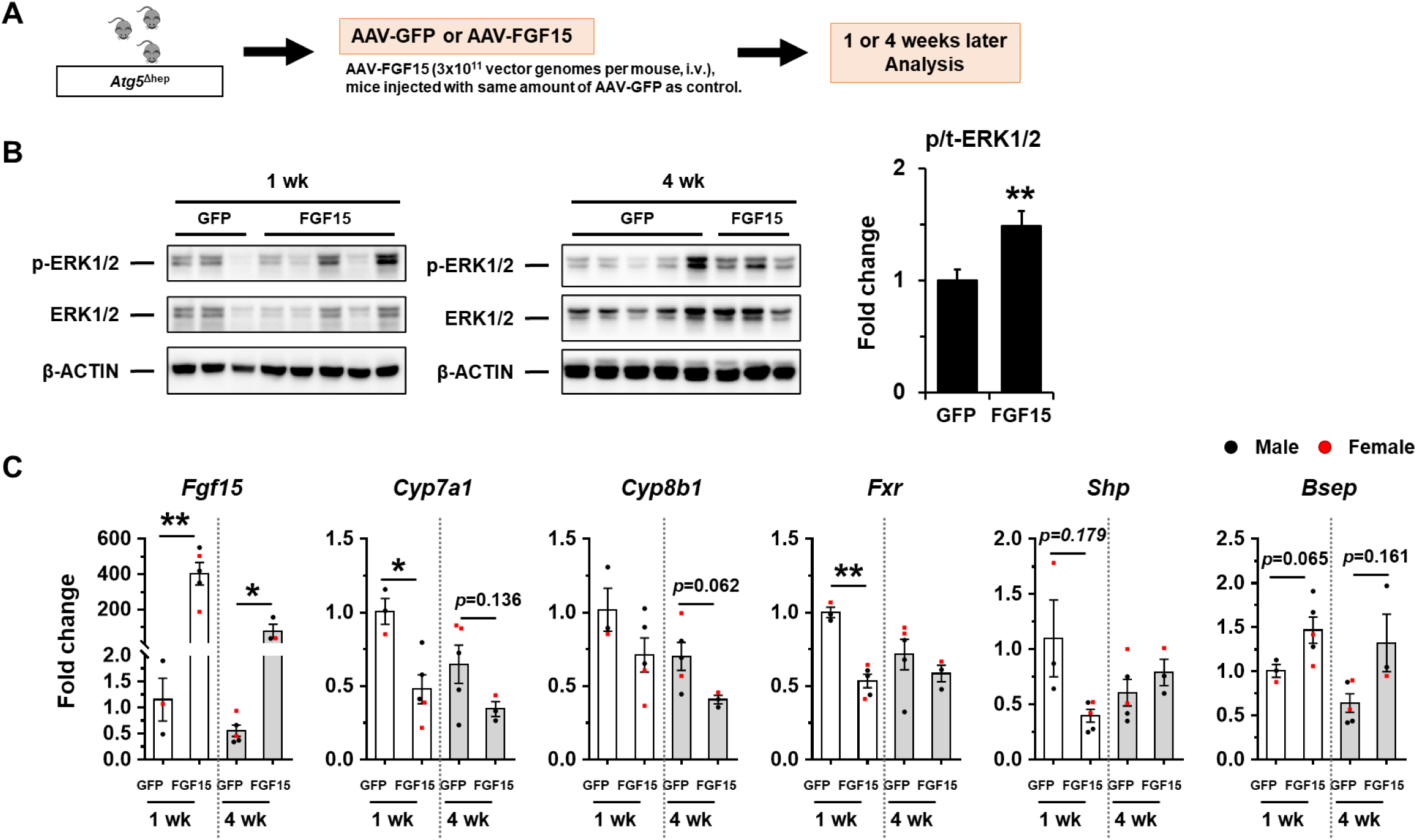
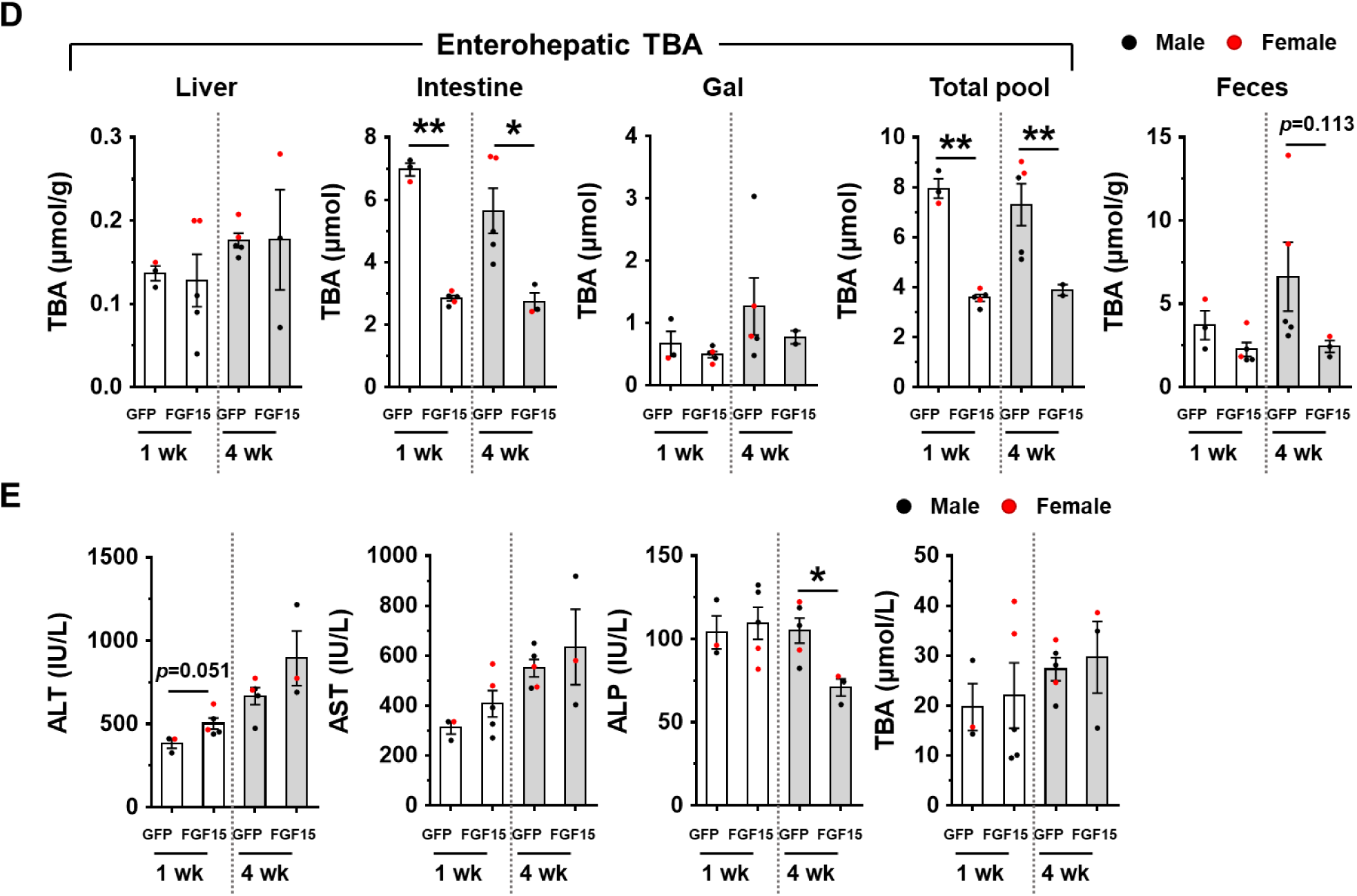
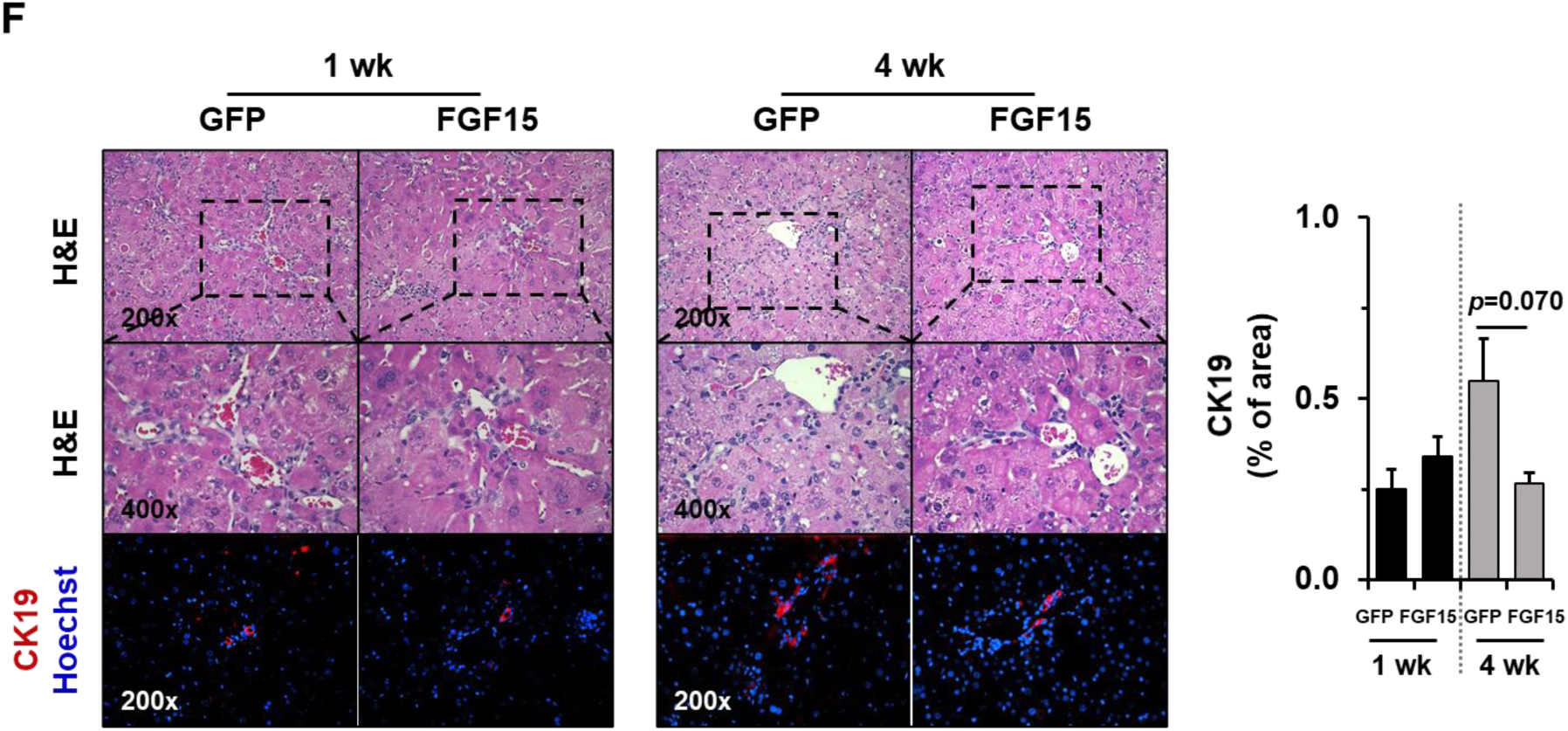
Overexpression of FGF15 attenuated pathological features in *Atg5*^*Δhep*^ livers. (**A**). Scheme of the FGF15 overexpression study in mouse livers. (**B**). Expression of ERK1/2 in the liver were analyzed by immunoblotting assay and quantified by densitometry. Phosphorylation levels of ERK1/2 were normalized to that of the total protein levels and expressed as fold change of GFP group (n=8/group). (**C**). Hepatic expression of indicated genes was analyzed by qRT-PCR (n=3-5/group). Data was expressed as fold change of GFP for the 1-week group. (**D**). TBA levels in the indicated compartments were measured (n=3-5/group). (**E**). The serum levels of ALT, AST, ALP, and TBA in mice following AAV-injection (n=3-5/group). (**F**). Liver sections were subjected to H&E, anti-CK19, or Masson’s trichrome staining. Percentage of positive area was quantified with ImageJ (n=3-5/group). Data were shown as means ± S.E. \**p*<0.05, \*\**p*<0.01. AAV, adeno-associated virus; ALT, alanine transaminase; ALP, alkaline phosphatase; AST, aspartate transaminase; *Bsep* (*Abcb11*), bile salt export pump; CK19, cytokeratin 19; *Cyp7a1*, cytochrome P450 7a1; *Cyp8b1*, cytochrome p450 8b1; ERK, extracellular-signal-regulated kinase; *Fgf15*, fibroblast growth factor 15; *Fxr*, farnesoid X receptor; *Shp*, small heterodimer partner; TBA, total bile acids.

Serum biochemistry analysis suggested FGF15 overexpression in the liver reduced ALP, but not ALT, AST, and TBA (Fig. 6E) in *Atg5*^Δhep^ mice, suggesting a potential improvement in biliary injury. Consistently, the H-E staining and CK19 staining showed a reduced ductular reaction (Fig. 6F). On the other hand, the parameters of fibrosis (Fig. S10) showed a variable improvement to FGF15 overexpression, suggesting that the short treatment had a minor impact on this process.

### Inhibition of FGFR4 aggravated liver injury in Atg5^Δhep^ mice

FGFR4 is the receptor that mediates FGF15 signaling in the liver [19]. In order to determine the role of FGF15-FGFR4 signaling in autophagy-deficient livers, we treated mice with Blu-9931 (BLU), a novel small molecular that selectively inhibits FGFR4 [30], to block FGFR4 activity (Fig. 7A).

**Fig. 7.**
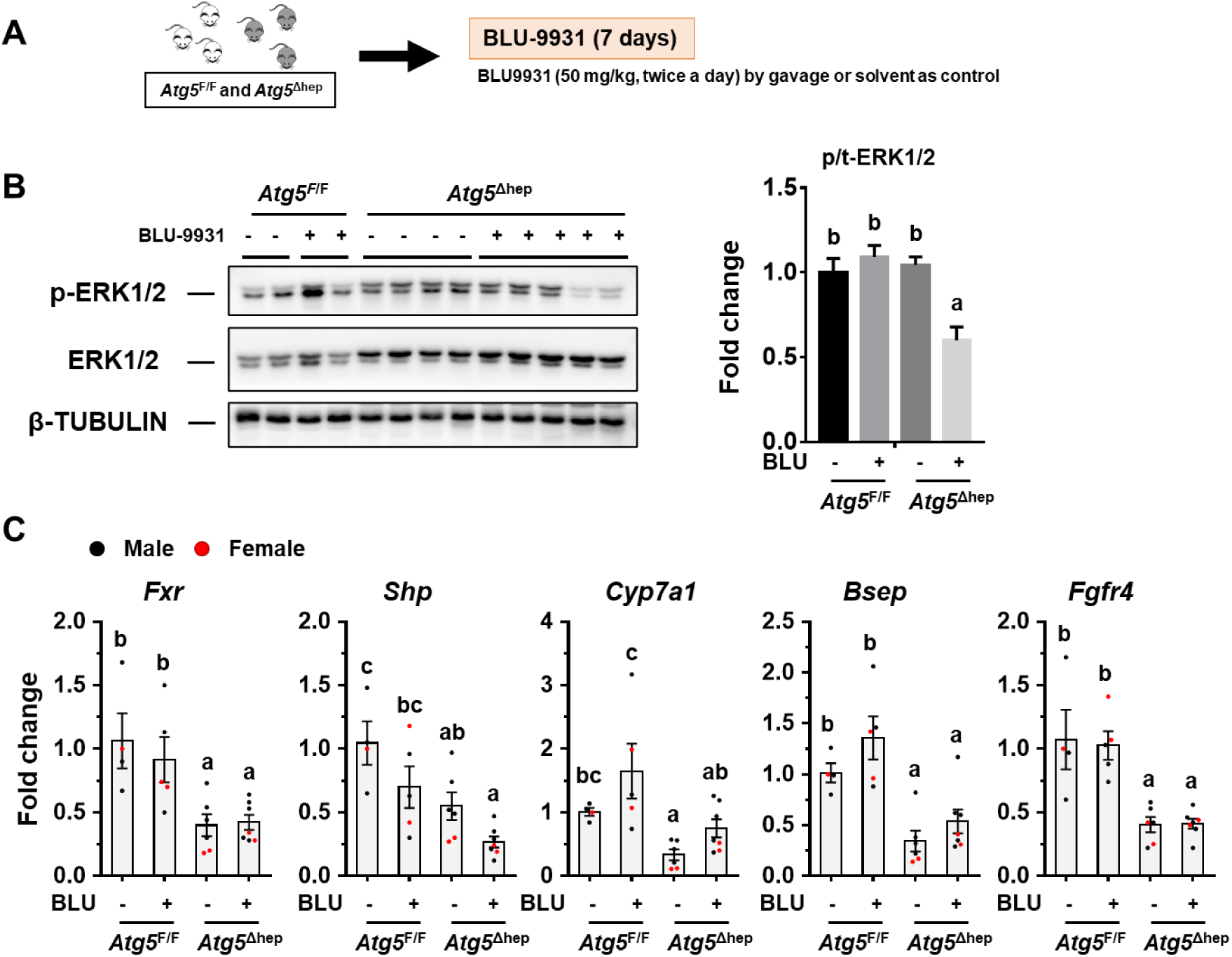
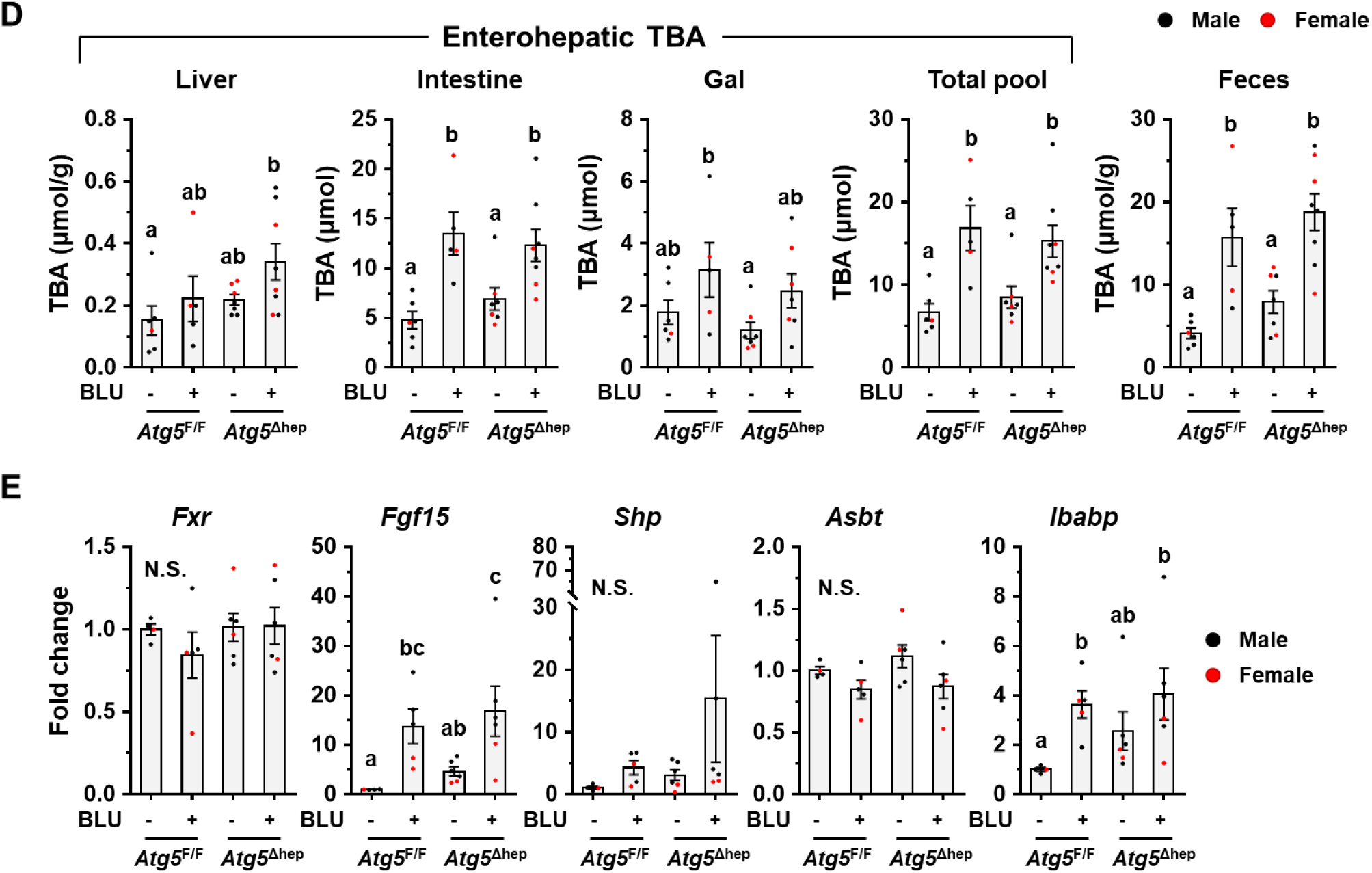
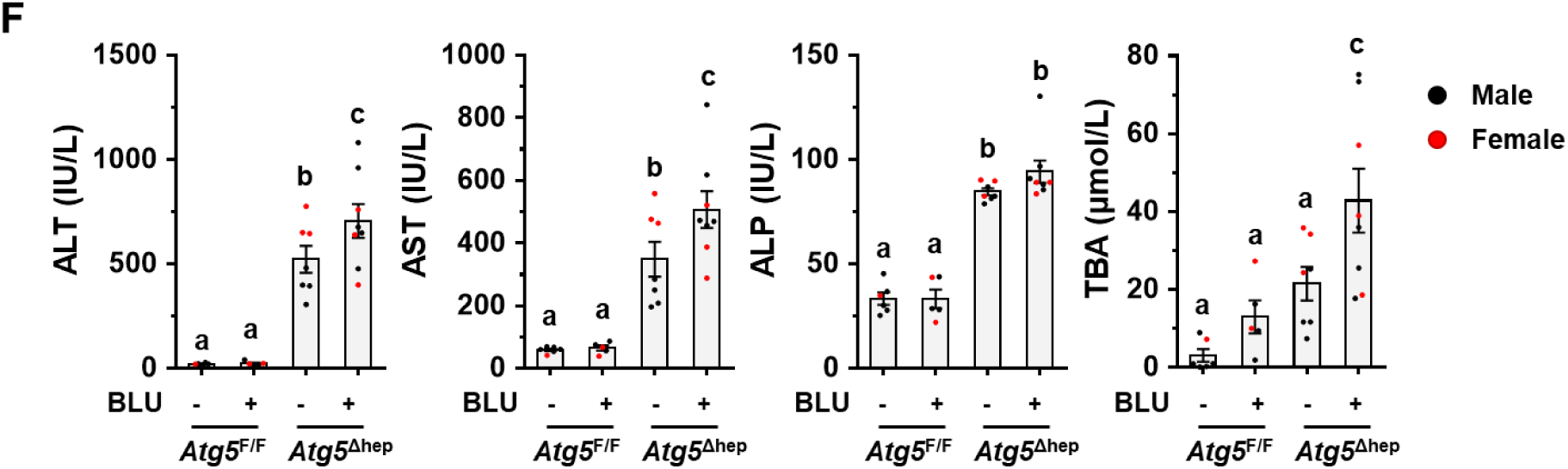
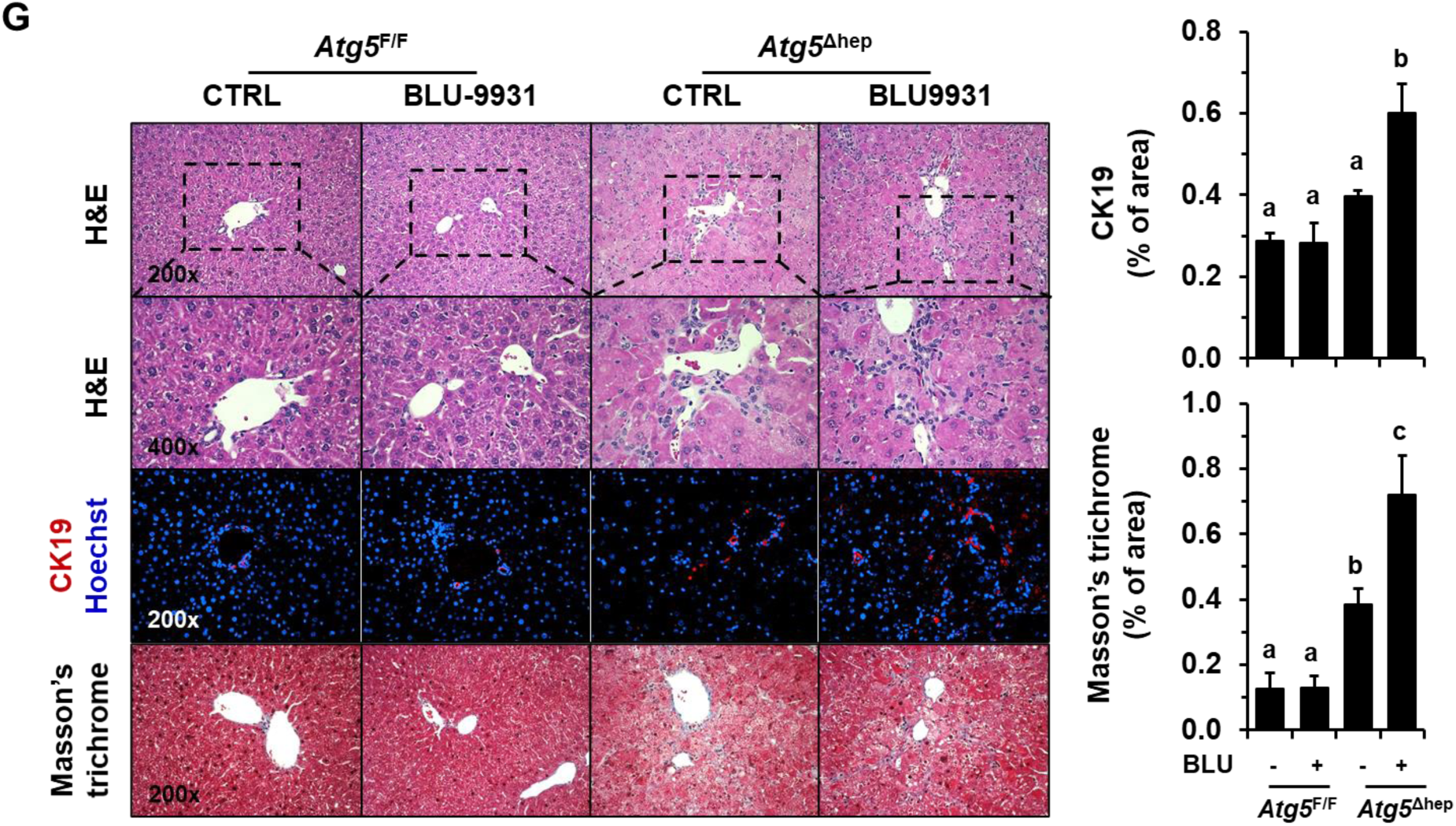
Inhibition of FGFR4 aggravated liver injury in *Atg5*^*Δhep*^ mice. (**A**). Scheme of, the treatment with FGFR4 inhibitor, Blu-9931 (BLU). Solvent (0.5% methylcellulose/1% Tween 80) was given as control. (**B**). The levels of ERK1/2 were examined by immunoblotting assay, and quantified by densitometry (n=3-5/group). The phosphorylation level of ERK1/2 was normalized to that of the total protein level and expressed as the fold change of *Atg5*^F/F^ control group. (**C**). Hepatic expression of indicated genes was analyzed by qRT-PCR (n=4-7/group). (**D**). TBA levels in indicated compartments were measured (n=6-8/group). (**E**). Ileal expression of indicated genes was analyzed by qRT-PCR (n=4-6/group). (**F**). The serum levels of ALT, AST, ALP, and TBA in mice following BLU-treatment (n=6-8/group). (**G**). Liver sections were subjected to H&E, anti-CK19, or Masson’s trichrome staining. Percentage of positive area was quantified with ImageJ (CK19 staining quantification, n=3-6/group; Masson’s trichrome staining quantification, n=4-5/group). Data were shown as means ± S.E. Groups with different letters had significant differences (*p<0.05*). N.S. indicates no statistical significance. ALT, alanine transaminase; ALP, alkaline phosphatase; AST, aspartate transaminase; *Asbt* (*Slc10a2*), apical sodium–bile acid transporter; *Bsep* (*Abcb11*), bile salt export pump; CK19, cytokeratin 19; *Cyp7a1*, cytochrome P450 7a1; ERK, extracellular-signal-regulated kinase; *Fgf15*, fibroblast growth factor 15; *Fgfr4*, fibroblast growth factor receptor 4; *Fxr*, farnesoid X receptor; *Ibabp*, ileal bile acid-binding protein; *Shp*, small heterodimer partner; TBA, total bile acids.

Phosphorylated ERK1/2 level was decreased in *Atg5*-deficient livers following BLU treatment (Fig. 7B), suggesting FGF15-FGFR4 signaling was suppressed in the liver. In both *Atg5*^F/F^ and *Atg5*^Δhep^ mice, BLU treatment reduced *Shp* expression whereas induced *Cyp7a1* expression in the liver (Fig. 7C). Consequently, the TBA pool and fecal TBA level were significantly increased following BLU treatment (Fig. 7D), indicating an increase of BA synthesis in the liver. Consistently, with the inhibition of hepatic FGFR4, ileal expression of *Fgf15* and *Ibabp* was remarkably induced following BLU treatment (Fig. 7E), which is possibly attributed to the increase of intestinal TBA.

Following BLU treatment, liver injury was enhanced in *Atg5*^Δhep^ mice as indicated by the significantly elevated serum levels of ALT, AST, and TBA (Fig. 7F). Notably, the male mice were more susceptible to BLU treatment for the serum enzyme activation (Fig. S11). We did not see significant pathological changes in livers of *Atg5*^F/F^ mice (Fig. 7G), suggesting that inhibition of FGFR4 was not toxic in healthy livers. However, the ductular reaction around the periportal areas was further enhanced in *Atg5*-deficient livers following BLU treatment as indicated by H&E staining and the positive areas of CK19 staining (Fig. 7G). Positive staining of Masson’s Trichrome were also significantly increased in *Atg5*-deficient livers following BLU treatment (Fig. 7G). Although we found a noticeable sexual disparity in serum enzyme changes, changes in histological studies were comparable between male and female *Atg5*^Δhep^ mice. Taken together, these results suggest that FGF15-FGFR4 signaling protects livers from further injury in *Atg5*-deficient mice.

## DISCUSSION

### Interaction between GM and hepatic autophagy deficiency and bile acid metabolism

In this study, we showed that autophagy deficiency in the liver led to the alteration of intestinal BA composition and GM with a significantly higher proportion of BA-metabolizing bacteria. Unexpectedly, ABX treatment increased enterohepatic level of BAs and exacerbated the pathology in autophagy-deficient livers. Together with other evidence, we demonstrate that enhanced activation of ileal FXR-FGF15 signaling, due to the effects of altered BA metabolism and GM, is accounted for the protection of the *autophagy*-deficient liver from further injury (Fig. 8). Therefore gut dysbiosis in liver diseases can be an adaptive response to mitigate the injury via a gut-liver signaling pathway.

**Fig. 8.**
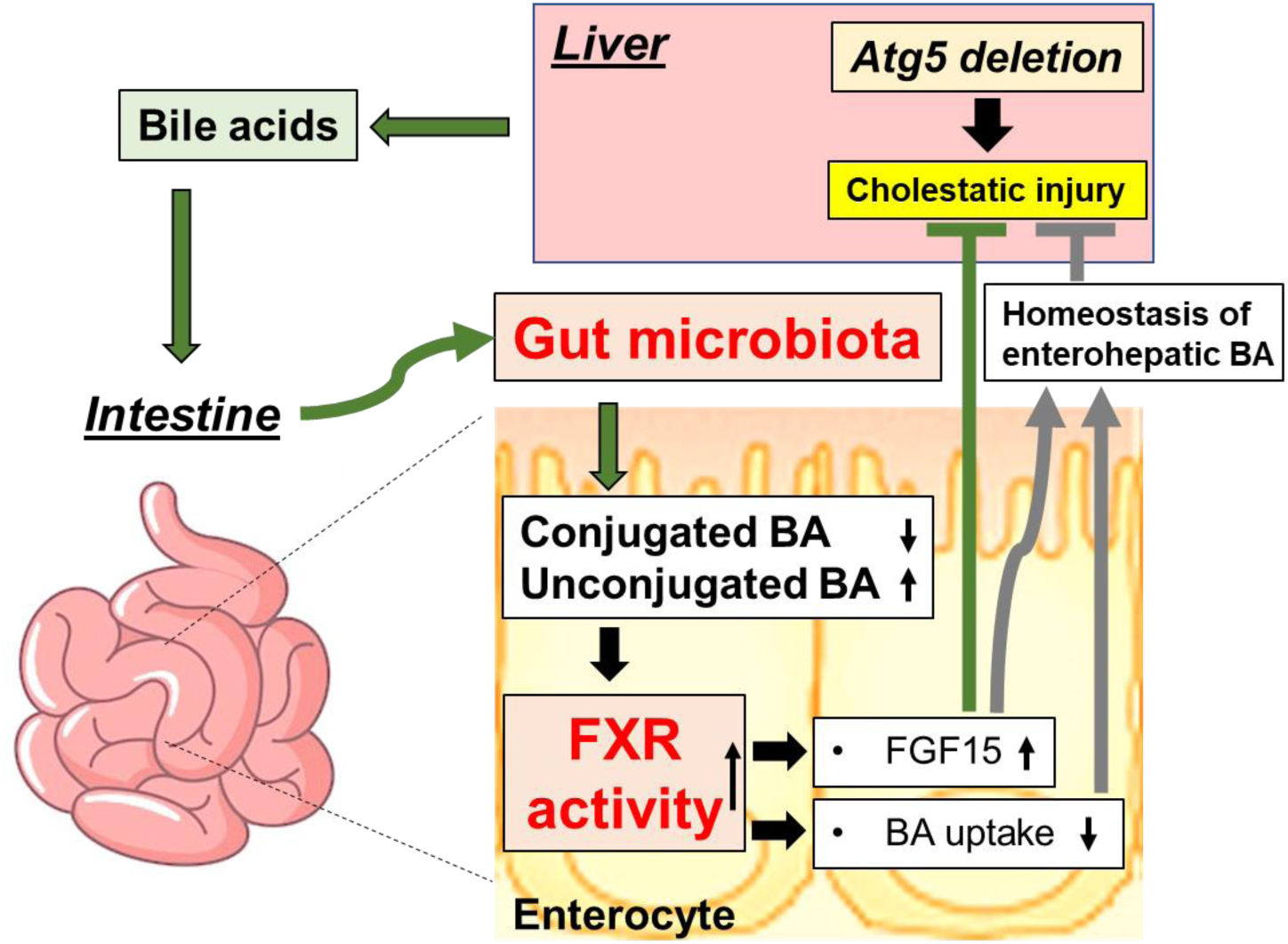
GM-dependent FXR-FGF15 signaling improves hepatic presentation in *Atg5*-deficient mice. *Atg5* deficiency in the liver impairs autophagy process and BA homeostasis, which causes liver injury accompanied with cholestasis. Disruption of hepatic BA homeostasis altered the composition of BAs in the intestine, and gut microbiota, which in turn affects BA composition in the intestine. The changes in intestinal BA composition activates ileal FXR, and consequently induces FGF15 expression and reduces BA uptake in the intestine, both of which can improve the homeostasis of enterohepatic BA. Importantly, FGF15 is beneficial to the improvement of the liver injury induced by autophagy deficiency. *Atg5*, autophagy-related gene 5; BA, bile acid; FGF15, fibroblast growth factor 15; FXR, farnesoid X receptor.

Autophagy deficiency in the liver causes hepatomegaly, chronic injury, and tumorigenesis [9, 10]. Mechanically, consistent activation of NRF2 by Sequestosome-1 (SQSTM1) is critical for pathological changes induced by hepatic *Atg5* or *Atg7*-deletion [9, 12, 13], yet how SQSTM1-NRF2 signaling leads to hepatocyte death remains unclear. Our previous study shows a compromised FXR activity and altered BA homeostasis in autophagy-deficient livers [11], in which activation of FXR in the liver can ameliorate *autophagy* deficiency-induced liver injury, suggesting that reduced hepatic FXR expression and disrupted BA homeostasis may at least partially contribute to injury in autophagy-deficient livers.

In this study we have defined a unique interaction between GM and liver injury in the context of autophagy deficiency, which affects BA metabolism and gut-liver signaling. Firstly, we show that dysfunction of BA metabolism induced by *Atg5* or *Atg7* deficiency in the liver alters GM and leads to a significant enrichment of BA-metabolizing bacteria with BSH activity. GM is critical for BA metabolism by deconjugation of BAs and conversion of primary BAs into secondary BAs [6]. Consistent with enriched BA-metabolizing bacteria, we observed a lower level of tauro-conjugated BAs but a higher level of unconjugated BAs in the intestine. However, levels of most secondary BAs in the intestine are lower in *Atg5*^Δhep^ intestines except deoxycholic acid (DCA). In mice, DCA is converted from TCA, whereas other secondary BAs are converted from TCDCA [31]. Interestingly, TCDCA level is decreased in *Atg5*^Δhep^ livers, suggesting a reduced capability of CDCA synthesis in these mice, which may contribute to the decreased levels of non-DCA secondary BAs in the intestine. Overall, the evidence here suggests that autophagy deficiency in the liver alters hepatic BA metabolism, which generates a different BA profile and favors the growth of a specific set of GMs.

Secondly, we found that altered GM maintains the enterohepatic BA level. We have reported cholestatic injury in autophagy-deficient livers with a significant increase of TBA levels in serum and livers [11]. Here, we not only confirmed our previous findings but also found that despite the increase of hepatic TBA level, the enterohepatic TBA pool is comparable between *Atg5*^F/F^ and *Atg5*^Δhep^ mice (Fig. 2F). Increased fecal elimination was found in *Atg5*^Δhep^ mice, which is reduced following ABX treatment, suggesting that GM maintained enterohepatic level of TBA in *Atg5*^Δhep^ mice by an increased BA excretion from the intestine. The mechanism of this increased excretion is not known, but could be related to a GM-dependent elevated ileal FXR activation, which needs to be further investigated in the future.

Thirdly, we present evidence that GM-mediated FXR activation in the ileum can induce FGF15 expression, thereby protecting mice from further liver damage caused by autophagy deficiency in the liver. Both the ABX and BAS treatments lead to a dramatic inhibition of ileal FXR activity and a significant decrease of FGF15 expression. Evidence from overexpression of FGF15 and inhibition of FGFR4 in the *Atg5*-deficient liver suggests that GM-mediated FGF15 expression at least partially protected *Atg5*^Δhep^ mice from further liver damage by a FGF15-FGFR4 feedback signaling pathway.

We previously found that *Atg7* deletion induced more severe pathological changes than *Atg5*-deletion in the liver, which leads to different response to alcohol treatment [32]. In this study, we also observed that ABX treatment enhanced liver injury in *Atg5*^Δhep^ but not obviously in *Atg7*^Δhep^ mice, despite increased BA pool and reduced ileal FXR activity following ABX treatment were observed in both *Atg5*^Δhep^ and *Atg7*^Δhep^ mice. It is possible that the protection effect from gut dysbiosis is overcome by additional hepatic phenotypes exerted by the more severe damage in the absence of *Atg7*, which sits on the upstream of Atg5 in the autophagy signaling pathway.

### Gut dysbiosis can be an adaptive response to liver injury

A number of studies had shown that gut dysbiosis contributes to the progress of liver disease and correction of the dysbiosis may improve pathological changes in the liver [33, 34, 35, 36, 37]. Among the detriment effects of GM, “invasion” of the liver by the product of GM due to increased gut permeability is thought to be the leading cause [4]. While the detrimental role of gut dysbiosis seems to be a widely recognized, there is also evidence indicating a beneficial impact of GM on acute liver injury in mice. Enrichment of intestinal *Lactobacillus* was found in mice with liver injury induced by acute concanavalin A (Con A) treatment, which can prevent further liver inflammation through activation of IL-22 production [38]. Furthermore, conflicting evidence supporting either a detrimental [39, 40] or beneficial [14, 41] effect of GM can be found for the liver injury in the ATP-binding cassette, sub-family B (MDR/TAP), member 4 (*Mdr2*) knockout (*Mdr2*^*-/-*^) mice.

In the present study, we have found GM is altered in *Atg5*^Δhep^ mice. Surprisingly, ABX treatment enhanced *Atg5* deficiency-induced liver injury, clearly indicating a protective role of GM in *Atg5*^Δhep^ mice. We further identified an increase of FXR activity and ileal FGF15 expression in *Atg5*^Δhep^ mice, which is associated with the altered intestinal BA composition and the dysbiosis status of GM. Our finding demonstrates that FGF15 can be a beneficial feedback signal from gut dysbiosis attributed to hepatic autophagy deficiency.

In mice, FGF15 is induced by FXR activation in the ileum, and its human ortholog is FGF19 [18]. FGF15/19 is required for the efficiency of SHP-mediated CYP7A1 repression and plays a critical role in repressing BA synthesis [18]. Conversely, decrease of intestinal level of BA by BA sequestrants can reduce ileal FGF15 expression in mice [26] and serum FGF19 levels in healthy humans [42]. Animal experiments have shown that FGF15 is essential for hepatic homeostasis, and overexpression of FGF15/19 in the liver has beneficial effects on multiple liver diseases, including sclerosing cholangitis [20], alcoholic fatty liver [22], and NAFLD [21, 29]. In addition to its functions in hepatic metabolism, FGF15 has also been shown to contribute to liver regeneration [23, 43, 44]. In human, circulating FGF19 has been found to be increased in patients with biliary cirrhosis [45] and in patients with alcoholic hepatitis [46], which is accompanied with inhibition of BA synthesis. Nevertheless, the function and cause of increased levels of circulating FGF19 remains unclear. Our current finding of the beneficial effects of FGF15-FGFR4 signaling in autophagy-deficient livers provides evidence that increase of FGF15/19 level in the setting of liver diseases can potentially be a protective mechanism via the gut-liver interaction.

Our study suggests that GM can adapt to metabolic changes in the liver, and consequently activate feedback signaling, like FXR-FGF15 signaling, in the gut to protect the liver from further damage. The present study also suggests cautions should be exercised in the use of antibiotics during specific liver diseases to avoid potential detrimental effects, not due to reduced hepatic drug metabolism, but due to disruption of beneficial gut-liver signaling.

## CONCLUSION

In summary, the findings in our studies indicate a primary liver disease can lead to alteration of GM, which then activates FXR-FGF15 feedback signaling through modulating the composition of intestinal BAs. Our results suggest that ABX treatment can exacerbate hepatic pathogenesis in *Atg5*-deficient livers by the reduction of FGF15 expression. Taken together, our present study demonstrates a protective role of gut dysbiosis in liver injury, which is associated with the FXR-FGF15 feedback signaling.

## ACKNOWLEDGEMENTS

This work was supported in part by the USA National Institutes of Health (NIH) grants DK116605 (to X.-M. Yin).

## Notes

### Competing Interest Statement

The authors have declared no competing interest.

