## Supplementary Material for "Gut dysbiosis protects against liver injury in autophagy deficient mice by FXR-FGF15 feedback signaling"

#### MATERIALS AND METHODS

##### *Animals and treatments*

*Atg5*$^{F/F}$ mice (*B6.129S-Atg5tm1Myok*) [1] and *Atg7*$^{F/F}$ mice [2] had been reported in previous studies. *Atg5*$^{\Delta hep}$ and *Atg7*$^{\Delta hep}$ mice were created by cross *Atg5*$^{F/F}$ or *Atg7*$^{F/F}$ with the Alb:Cre transgenic mice (The Jackson Laboratory, Bar Harbor, ME), respectively. Mice were maintained on a 12-hour dark/12-hour light cycle with free access to food and water. Both male and female mice were used in the studies if not further addressed. Age and sex matched mice were randomly assigned to treatment or control group. For antibiotics treatment, mice (5-6 weeks old) were given antibiotics (0.5g/L neomycin sulfate and 1g/L ampicillin) in daily drinking water for 6 weeks. For bile acid sequestrant treatment, cholestyramine resin (2 g/kg) was mixed in water and given to 8-12 weeks old mice twice a day by oral gavage for 5 days. Control groups were given a same volume of water. For fexaramine treatment, fexaramine was dissolved in DMSO and further diluted in corn oil according to manufacturer’s protocol. Mice (8-12 weeks old) were given 50 mg/kg fexaramine daily by oral gavage for 7 days. Control mice were given the same volume of corn oil containing the same amount of DMSO. For adeno-associated virus (AAV)-mediated overexpression of FGF15, *Atg5*$^{\Delta hep}$ mice (6-8 weeks old) were given $3x10^{11}$ vector genomes of AAV-FGF15 per mouse by *i.v.* injection. Control mice were given AAV-GFP. Mice were euthanized for further analysis 1 or 4 weeks later. For FGFR4 inhibitor treatment, Blu-9931 was dissolved in 0.5% methylcellulose/1% Tween 80 solution according to a previous study [3]. Mice (8-12 weeks old) were given 50 mg/kg Blu-9931 twice per-day by oral gavage for 7 days. Control mice were given the same volume of solvent. All animal experiments were approved by the Institutional Animal Care and Use Committee (IACUC) of Indiana University.

##### *Antibodies and chemicals*

Antibodies and PCR primers used in this study are listed in Supplemental Table 1 and Supplemental Table 2, respectively.

***Fecal 16S rRNA sequencing of gut microbial communities***

Mice were 6-26 weeks old when fecal samples were collected. For *Atg5*$^{\Delta hep}$ mice, fecal samples were collected from *Atg5*$^{F/F}$ and *Atg5*$^{\Delta hep}$ mice at 8- or 16-weeks old. For *Atg7*$^{\Delta hep}$ mice, fecal samples were collected from floxed *Atg7* mice (heterozygous or homozygous, *Atg7*$^{WT}$) and *Atg7*$^{\Delta hep}$ mice. Both male and female mice were used and equally distributed in different genotypes. Fecal DNA was extracted from frozen fecal samples using the E.Z.N.A. Stool DNA Kit (Omega Bio-Tek, Inc., Norcross, GA, USA). All DNA samples were stored at -80°C before sequencing, which was performed by SeqMatic LLC (Fremont, CA, USA) using Illumina sequencing libraries. FASTQ data was processed using the Qiime pipeline on Illumina's BaseSpace servers.

For 16S sequencing analysis, relative abundance of each bacteria was calculated. Principle coordinates analysis (PCoA) was performed using multidimensional scaling function based on relative abundance at species level using SPSS for Windows 17.0 Software (SPSS, Inc., Chicago, IL, USA). Heatmaps were generated were generated using Morpheus (https://software.broadinstitute.org/morpheus) and values in the heatmap were mapped to colors using the minimum and maximum of each row independently. The hierarchical cluster of each heatmap was constructed using one minus Pearson correlation method.

***Serum biochemistry analysis***

Serum levels of alanine aminotransferase, aspartate aminotransferase, and alkaline phosphatase were measured using kits from Pointe Scientific (Canton, MI) according to the manufacturer's protocol. Serum total bile acids were measured using the total bile acids assay kit from Diazyme Laboratories, Inc (Poway, CA).

***Tissue and fecal TBA content analysis***

Sample preparation and bile acids quantification were performed as in previous studies with modifications [4]. For the liver tissue, samples (100 mg) were homogenized in 1 mL of 90% ethanol and incubated at 55°C overnight. The lysates were centrifuged at 10,000 rpm for 10 min. Supernatant was used to measure bile acids concentration. The whole intestine (with its content) was homogenized in 5ml of water, and was incubate at 55°C overnight after addition of 45 ml ethanol. One milliliter of the lysates was removed for centrifugation at 10,000 rpm for 10 min. An aliquot of the supernatant was diluted 5 times and measured for bile acids concentration. For bile acids in the gallbladder, the entire organ was put in 1 mL of 90% ethanol. After the gallbladder was cut to release the bile, the samples were Incubated at 55°C overnight. The lysates were centrifuged at 10,000 rpm for 10 min. An aliquot of the supernatant was diluted 50 times to measure the bile acids concentration. For fecal bile acids measurement, overnight-dried fecal samples (150-250 mg) were admixed with 1 mL of 95% EtOH and incubated at 55°C overnight. The lysates were centrifuged at 10,000 rpm for 10 min. An aliquot of the supernatant was diluted 10 times for bile acids quantification. The bile acids concentration of each diluted supernatants was measured using the total bile acids assay kit from Diazyme Laboratories, Inc (Poway, CA).

***Liver and intestinal bile acids composition analysis***

Liver and intestinal samples from male mice were used for bile acids profile analysis. Bile acids analysis was using a Biocrates® Bile Acids Kit (Biocrates Life Science AG, Innsbruck, Austria). Samples were prepared according to the manufacturer's protocol with modifications. At least 30 mg of liver tissue were homogenized in 3-fold volume of extraction buffer (ethanol/phosphate buffer, 85:15 v/v). For intestine samples, they were homogenized in 3 mL of PBS, and then 17 mL of 100% ethanol were added to extract bile acids. All samples were probe-sonicated for 3 bursts with 10 seconds each. Samples were chilled in an ice bath for at least 60 seconds between bursts. Homogenized samples were stored at -80°C. Further bile acids extraction and profile analysis were performed according to manufacturer's protocol in Center for Genomic and Computational Biology, Duke University. Sample pool quality controls (SPQC) were created using equal volumes of all liver and intestine samples respectively. Bile acids were identified using Biocrates Me*tIDQ*™ software. A Principal Components Analysis (PCA) was performed for bile acids using JMP® Pro v14.0 software (SAS, Cary, NC).

For bile acids composition analysis, results of analytes with more than 40% missing values were removed before statistical analysis. Missing values of the rest results were then replaced with the limit of detection (LOD). Results from liver samples were calculated as nmol/g or percentage of total bile acids level, and were calculated as nmol/whole intestine or percentage of total bile acids level from intestine samples. Heatmaps were generated were generated using Morpheus (https://software.broadinstitute.org/morpheus) and values in the heatmap were mapped to colors using the minimum and maximum of each row independently. The hierarchical cluster of each heatmap was constructed using one minus Pearson correlation method.

##### *Immunoblotting analysis*

The liver samples were homogenized in the radio immunoprecipitation assay buffer (RIPA) containing a protease cocktail and phosphatase inhibitors. Supernatant was collected after centrifugation at 12,000×rpm for 12 min. Protein concentration was determined using BCA protein assay kit (Thermo Fisher Scientific-Pierce). The proteins were separated on a sodium dodecyl sulfate polyacrylamide gel electrophoresis (SDS-PAGE). Proteins were transferred onto polyvinylidene fluoride (PVDF) membranes, which were then blocked with 5% BSA or 5% skim milk for 1 hour at room temperature. Membranes were incubated with the appropriate primary antibody overnight and then washed with TBST (Tris-buffered saline, 0.1% Tween 20) before being incubated with the horseradish peroxidase-coupled secondary antibodies at room temperature. Protein bands were detected using enhanced chemiluminescence (ECL) kit (Thermo Fisher Scientific-Pierce). The images were taken digitally with a BioRad ChemiDoc Image System (BioRad, Hercules, CA). Densitometry was measured using the companion software, and the values were normalized to that of β-TUBULIN, β-ACTIN or GAPDH, which were then converted to fold change of the control.

##### *RNA isolation and quantitative real-time PCR analysis*

Total RNA was prepared from liver samples using GeneTET RNA purification kit (Thermo Fisher Scientific, Grand Island, NY) according to manufacturer's protocols. Complementary DNA (cDNA) was synthesized using Oligo dT primers and an M-MLV reverse transcriptase system (Life Technologies-Thermo Fisher Scientific). Quantitative real-time PCR analysis (qRT-PCR) was performed on a QuantStudio 3 Real-Time PCR System (Life Technologies-Applied Biosystems) using SYBR Green master mixes (Life Technologies-Applied Biosystems). All qRT-PCR results were normalized to the level of β-actin and the gene expression was calculated using the $2^{-\Delta\Delta Ct}$ method.

##### *Histological study*

Liver or ileum samples were rinsed with PBS, and fixed in 10% formalin overnight. Samples were further fixed in 70% ethanol and processed as paraffin-embedded blocks. The paraffin-embedded liver tissues were sectioned and stained with hematoxylin and eosin (H&E), or Mason's Trichrome C. The paraffin-embedded ileum tissues were sectioned and performed immunohistochemical staining with FGF15 antibody. Photomicrographs were taken using a Nikon Eclipse E200 light microscope equipped with a SPOT RT Slider color digital camera (Diagnostic Instruments, Inc, Sterling Heights, MI). Area with positive Mason's Trichrome C staining was quantified using ImageJ software (NIH), at least 4 random fields of each section from each mouse liver were used for quantification.

##### *Immunofluorescence microscopy*

Paraffin sections were subjected to antigen retrieval treatment using the Citrate buffer (0.01M, pH 6.0) after deparaffinization. Slides were blocked with 5% goat serum in PBS containing 0.1% Triton X (PBS-Tx) for 1 hour and then incubated with primary antibodies diluted in 1% bovine serum albumin (BSA)/PBS-Tx overnight at 4°C. Sections were washed with PBS, following by incubation with fluorochrome-conjugated secondary antibodies. Hoechst 33342 was used for nucleus staining. Images were obtained using a Nikon Eclipse TE 200 epi-immunofluorescence microscope and the companion NIS-Elements AR3.2 software. Quantification was performed using ImageJ software (NIH), at least 4 random fields of each section from each mouse liver were analyzed.

##### *Statistical analysis*

The 16S sequencing data were represented as median with interquartile range. All the other data were represented as means with standard errors (mean ± S.E.). For 16S sequencing data, Mann-Whitney test was performed to identify bacteria with significantly different proportions between *Atg5*$^{\Delta hep}$ and sex- and age-matched *Atg5*$^{F/F}$ mice. For all the other data, to determine statistical significance, student's *t*-test was used to determine differences between two groups. Differences among more than two treatment groups were determined using one-way analysis of variance (ANOVA) followed by Duncan's post-hoc test. Results were considered statistically significant for *p*-value < 0.05. Statistical analyses were performed using SPSS for Windows 17.0 Software (SPSS, Inc., Chicago, IL, USA).

**REFERENCES** (limited to those critical and relevant to the manuscript, **around 50**)


1 Takamura A, Komatsu M, Hara T, Sakamoto A, Kishi C, Waguri S*, et al.* Autophagy-deficient mice develop multiple liver tumors. Genes Dev 2011;**25**:795-800.

2 Komatsu M, Kurokawa H, Waguri S, Taguchi K, Kobayashi A, Ichimura Y*, et al.* The selective autophagy substrate p62 activates the stress responsive transcription factor Nrf2 through inactivation of Keap1. Nat Cell Biol 2010;**12**:213-23.

3 Hagel M, Miduturu C, Sheets M, Rubin N, Weng W, Stransky N*, et al.* First Selective Small Molecule Inhibitor of FGFR4 for the Treatment of Hepatocellular Carcinomas with an Activated FGFR4 Signaling Pathway. Cancer Discov 2015;**5**:424-37.

4 Pathak P, Xie C, Nichols RG, Ferrell JM, Boehme S, Krausz KW*, et al.* Intestine farnesoid X receptor agonist and the gut microbiota activate G-protein bile acid receptor-1 signaling to improve metabolism. Hepatology 2018;**68**:1574-88.

**ABBREVIATIONS:**

27-HOC, 27-hydroxycholesterol

7α-HOC, 7α-hydroxycholesterol

ABX, antibiotics

AKR1D1, aldo-keto reductase family 1 member d1

ALP, alkaline phosphatase

ALT, alanine transaminase

ASBT (SLC10A2), apical sodium–bile acid transporter

AST, aspartate transaminase

ATG, autophagy-related gene

BAAT, bile acid-coa:amino acid n-acyltransferase

BA, bile acid

BAS, bile acid sequestrants

BLU, blu-9931

BSEP (ABCB11), bile salt export pump

BSH, bile salt hydrolase

CA, cholic acid

CDCA, chenodeoxycholic acid

CK19, cytokeratin 19

CYP27A1, cytochrome p450 27a1

CYP2C70, cytochrome p450 2c70

CYP7A1, cytochrome p450 7a1

CYP7B1, cytochrome p450 7b1

CYP8B1, cytochrome p450 8b1

DCA, deoxycholic acid

ERK, extracellular-signal-regulated kinase

FEX, fexaramine

FGF15, fibroblast growth factor 15

FGFR4, fibroblast growth factor receptor 4

FXR, farnesoid x receptor

GAPDH, glyceraldehyde-3-phosphate dehydrogenase

GM, gut microbiota

H&E, haemotoxylin and eosin

IBABP, ileal bile acid-binding protein

LC3B, microtubule associated protein 1 light chain 3 beta

MAMP, microbial-associated molecular pattern

MCA, muricholic acid

MDR1A (ABCB1), multidrug resistance protein 1a

MRP2 (ABCC2), multidrug resistance-associated protein 2

MRP3 (ABCC3), multidrug resistance-associated protein 3

MRP4 (ABCC4), multidrug resistance-associated protein 4

NAFLD, nonalcoholic fatty liver disease

NASH, nonalcoholic steatohepatitis

NQO1, nad(p)h quinone dehydrogenase 1

NRF2, nuclear factor erythroid 2-related factor 2

NTCP (SLC10A1), Na/Taurocholate cotransporting polypeptide

OATP1 (SLCO1A1), organic anion-transporting polypeptide 1

OST-A (SLC51A), organic solute transporter subunit α

OST-B (SLC51B), organic solute transporter subunit β

SHP, small heterodimer partner

SQSTM1, sequestosome-1

TCA, taurocholic acid

TCDCA, taurochenodeoxycholic acid

TDCA, taurodeoxycholic acid

TLCA, taurolithocholic acid

TMCA, tauromuricholic acid

TUDCA, tauroursodeoxycholic acid

#### Table S1 Antibody list

| Antibody name | Company | Catalog # | Host |
|---|---|---|---|
| **ATG12** | Cell Signaling | 2011 | rabbit |
| **β-ACTIN** | Cell Signaling | 3700 | mouse |
| **CK19** | DSHB | TROMA-III | rat |
| **Phospho-ERK1/2 (Thr202/Tyr204)** | Cell Signaling | 4370 | rabbit |
| **ERK1/2** | Cell Signaling | 9102 | rabbit |
| **FGF15** | Santa Cruz | sc-514647 | mouse |
| **GAPDH** | Novus biologicals | NB 300-221 | mouse |
| **LC3B** | Sigma | L7543 | rabbit |
| **NQO1** | Abcam | ab34173 | rabbit |
| **P62/SQSTM1** | Abnova | H00008878-M01 | mouse |
| **α-SMA** | Thermo | PA5-19465 | rabbit |
| **β-TUBULIN** | Cell Signaling | 86298 | mouse |

#### Table S2 Primer list

| Gene name | Sequence (Forward) | Sequence (Reverse) |
|---|---|---|
| ***Actin*** | 5’-ACTATTGGCAACGAGCGGTT-3’ | 5’-CAGGATTCCATACCCAAGAAGGA-3’ |
| ***Akr1d1*** | 5’-CTCATTGGGCTTGGAACCTA-3’ | 5’-CATTGATGGGACATGCTCTG-3’ |
| ***Asbt (Slc10a2)*** | 5’-CGACATGGACCTCAGTGTTAG-3’ | 5’-CAACCCACATCTTGGTGTAGA-3’ |
| ***Baat*** | 5’-GTCCTTTTCCAGGGGTCATT -3’ | 5’-CCAGAGCTAAGGTGGCAAAG -3’ |
| ***Bsep (Abcb11)*** | 5’-CTGCCAAGGATGCTAATGCA-3’ | 5’-CGATGGCTACCCTTTGCTTCT-3’ |
| ***Cyp27a1*** | 5’-GCCTCACCTATGGGATCTTCA-3’ | 5’-TCAAAGCCTGACGCAGATG-3’ |
| ***Cyp7a1*** | 5’-AACAACCTGCCAGTACTAGATAGC-3’ | 5’-GTGTAGAGTGAAGTCCTCCTTAGC-3’ |
| ***Cyp7b1*** | 5’-CAGCTATGTTCTGGGCAATG-3’ | 5’-TCGGATGATGCTGGAGTATG-3’ |
| ***Cyp8b1*** | 5’-AGTACACATGGACCCCGACATC-3’ | 5’-GGGTGCCATCCGGGTTGAG-3’ |
| ***Fgfr4*** | 5’-CTGCCAGAGGAAGACCTCAC-3’ | 5’-GTAGTGGCCACGGATGACTT-3’ |
| ***Fgf15*** | 5’-ATGGCGAGAAAGTGGAACGG -3’ | 5’-CTGACACAGACTGGGATTGCT -3’ |
| ***Fxr (Nr1h4)*** | 5’-GGCCTCTGGGTACCACTACA-3’ | 5’-TGTACACGGCGTTCTTGGTA-3’ |
| ***Gstm1*** | 5’-ACTTGATTGATGGGGCTCAC-3’ | 5’-TCTCCAAAATGTCCACACGA-3’ |
| ***Ibabp*** | 5’-CCCCAACTATCACCAGACTTC-3’ | 5’-ACATCCCCGATGGTGGAGAT-3’ |
| ***Mdr1a (Abcb1)*** | 5’-AAAGGCTCTACGACCCCCTA-3’ | 5’-CCTGACTCACCACACCAATG-3’ |
| ***Mrp2 (Abcc2)*** | 5’-GCACTGTAGGCTCTGGGAAG-3’ | 5’-TGCTGAGGGACGTAGGCTAT-3’ |
| ***Mrp3 (Abcc3)*** | 5’-GGACTTCCAGTGCTCAGAGG-3’ | 5’-AGCTGTGGCCTCGTCTAAAA-3’ |
| ***Mrp4 (Abcc4)*** | 5’-TGTTTGATGCACACCAGGAT-3’ | 5’-GACAAACATGGCACAGATGG-3’ |
| ***Nqo1*** | 5’-GCACTGATCGTACTGGCTCA-3’ | 5’-CATGGCATAGAGGTCCGACT-3’ |
| ***Ntcp (Slc10a1)*** | 5’-CACCATGGAGTTCAGCAAGA-3’ | 5’-CCAGAAGGAAAGCACTGAGG-3’ |
| ***Ost-α (Slc51A)*** | 5’-GTCTCAAGTGATGAACTGCCA-3’ | 5’-TTGAGTGCTGAGTCCAGGTC-3’ |
| ***Ost-β (Slc51B)*** | 5’-GTATTTTCGTGCAGAAGATGCG-3’ | 5’-TTTCTGTTTGCCAGGATGCTC-3’ |
| ***Oatp1 (Slco1a1)*** | 5’-ATCCAGTGTGTGGGGACAAT-3’ | 5’-GCAGCTGCAATTTTGAAACA-3’ |
| ***Shp*** | 5’-CTGGTTGAGCGCCTGAGAC-3’ | 5’-CTGCCTGGATGCCCTTTATC -3’ |

**Fig. S1 Liver-specific deletion of *Atg7* caused gut dysbiosis.** Shannon species diversity (**A**) and the number of species identified (**B**) were analyzed in each group of mice. (**C**). Bacteria with disproportionated representation in *Atg7*$^{\Delta hep}$ mice at genus level. Heatmap was generated and values in the heatmap were mapped to colors using the minimum and maximum of each row independently. The hierarchical cluster of different genus was constructed using one minus Pearson correlation method. Male and female mice were 6-26 weeks old when fecal samples were collected (n=5). Data were shown as median with interquartile range. Mann-Whitney analysis did not show statistical significance between the groups.

**Fig.S2 Antibiotics treatment did not affect NRF2 activity in *Atg5* deficiency livers.** (**A**). Immunoblotting analysis of hepatic samples. (**B**). The mRNA level of *Nqo1* and *Gstm1* was determined by qRT-PCR (n=7-9). (**C**). Average daily water consumption per mouse was determined. Data were shown as means ± S.E. Groups with different letters had significant differences ($p<0.05$). GAPDH, glyceraldehyde-3-phosphate dehydrogenase; *Gstm1*, glutathione s-transferase mu 1; LC3B, microtubule associated protein 1 light chain 3 beta; *Nqo1*, NAD(P)H quinone dehydrogenase 1; SQSTM1, sequestosome-1.

**Fig.S3 ABX treatment suppressed ileal FXR activity and altered enterohepatic TBA levels in *Atg7*$^{\Delta hep}$ mice.** (**A**). The TBA levels in the indicated compartments were measured (n=4-6). (**B**). The serum levels of ALT, AST, ALP, and TBA in *Atg7*$^{\Delta hep}$ mice following ABX-treatment (n=4-6). (**C**). Liver weight and gallbladder (Gal) weight were shown as a percentage of body weight (n=4-6). Data were shown as means ± S.E. Groups with different letters had significant differences ($p<0.05$). ALT, alanine transaminase; ALP, alkaline phosphatase; AST, aspartate transaminase; TBA, total bile acids.

**Fig.S4 Alterations in the proportion of BA-metabolizing bacteria in *Atg7*$^{\Delta hep}$ mice.** (**A**). The proportion of bacteria with bile salt hydrolase and/or 7α/β-dehydroxylation activity at the *Genus* level in floxed *Atg7* or in *Atg7*$^{\Delta \mathrm{hep}}$ mice. Mice were 6-26 weeks old. Both male and female mice were included. (**B**). The bacterial species with bile salt hydrolase and/or 7α/β-dehydroxylation activity which were disproportionated in *Atg7*$^{\Delta hep}$ mice. Data were shown as median with interquartile range, n=5. Mann-Whitney analysis, *$p$<0.05, **$p$<0.01.

**Fig.S5 Hepatic autophagy deficiency caused changes in the proportion of BA-metabolizing bacteria.** (**A**). The Venn diagram shows the number of bacterial species with bile salt hydrolase and/or 7α/β-dehydroxylation activity, which were disproportionated in *Atg5*$^{\Delta hep}$ mice compared to the sex- and age- matched *Atg5*$^{F/F}$ mice. (**B**). Disproportionated bacteria at the *Species* level were segregated based on the gender of the mouse. Heatmap was generated and values in the heatmap were mapped to colors using the minimum and maximum of each row independently. The hierarchical cluster of different species was constructed using one minus Pearson correlation method.

**Fig.S6 The BA composition in the liver following ABX treatment.** (**A**). Principal components analysis of BAs in the liver (log2-scaled µM). (**B**). Heatmap was generated and values in the heatmap were mapped to colors using the minimum and maximum of each row independently. The hierarchical cluster of different BAs was constructed using one minus Pearson correlation method. (**C-D**). Hepatic levels of unconjugated bile acids (**C**), and Hepatic levels of tauro-conjugated bile acids (**D**) in male mice (n=5). Data were shown as means ± S.E. Groups with different letters had significant differences ($p$<0.05). CA, cholic acid; CDCA, chenodeoxycholic acid; DCA, deoxycholic acid; HDCA, hyodeoxycholic acid; LCA, lithocholic acid; MCA,muricholic acid; TCA, taurocholic acid; TCDCA, taurochenodeoxycholic acid; TDCA, taurodeoxycholic acid; TLCA, taurolithocholic acid; TMCA, tauromuricholic acid; TUDCA, tauroursodeoxycholic acid; UDCA, ursodeoxycholic acid.

**Fig.S7 ABX treatment suppressed ileal FXR activity and reduced *Fgf15* expression in *Atg7*$^{\Delta hep}$ mice.** Ileal expression of indicated genes was analyzed by qRT-PCR (n=4-6). Data were shown as means ± S.E. Groups with different letters had significant differences ($p$<0.05). *Asbt* (*Slc10a2*), apical sodium–bile acid transporter; *Fgf15*, fibroblast growth factor 15; *Fxr*, farnesoid X receptor; *Ibabp*, ileal bile acid-binding protein; *Shp*, small heterodimer partner.

**Fig.S8 The mRNA level of hepatic genes related to BA metabolism following ABX treatment.** (**A**). Expression of bile acid metabolism-related genes was analyzed by qRT-PCR in the liver (n=6-9). (**B**). Hepatic expression of BA transporters was analyzed by qRT-PCR (n=6-9). Data were shown as means ± S.E. Groups with different letters had significant differences. Akr1d1, aldo-keto reductase family 1 member d1; Baat, bile acid-coa:amino acid n-acyltransferase; *Bsep* (*Abcb11*), bile salt export pump; *Cyp7b1*, cytochrome P450 7b1; *Cyp8b1*, cytochrome P450 8b1; *Cyp27a1*, cytochrome P450 27a1; *Fgfr4*, fibroblast growth factor receptor 4; *Fxr*, farnesoid X receptor; *Mdr1a* (*Abcb1*), multidrug resistance protein 1; *Mrp2* (*Abcc2*), multidrug resistance-associated protein 2; *Mrp3* (*Abcc3*), multidrug resistance-associated protein 3; *Mrp4* (*Abcc4*), multidrug resistance-associated protein 4; *Ntcp* (*Slc10a1*), Na/Taurocholate cotransporting polypeptide; *Oatp1* (*Slco1a1*), organic anion-transporting polypeptide 1; *Ost-α* (*Slc51A*), organic solute transporter subunit α; *Ost-β* (*Slc51B*), organic solute transporter subunit β; *Shp*, small heterodimer partner.

**Fig.S9 Intestine-specific FXR agonist activated ileal FXR in *Atg5*$^{F/F}$ but not in *Atg5*$^{\Delta hep}$ mice.** (**A**). Scheme of the study using an intestine-specific FXR agonist, fexaramine (FEX). Solvent (1% DMSO in corn oil) was given as the control. (**B**). The serum levels of ALT, AST, ALP, and TBA in mice following FEX-treatment (n=5-8). (**C**). Ileal expression of indicated genes was analyzed by qRT-PCR (n=5). (**D**). TBA levels in the indicated compartments were measured (n=5-8). Data were shown as means ± S.E. Groups with different letters had significant differences ($p<0.05$). ALT, alanine transaminase; ALP, alkaline phosphatase; AST, aspartate transaminase; *Asbt* (*Slc10a2*), apical sodium–bile acid transporter; *Fgf15*, fibroblast growth factor 15; *Fxr*, farnesoid X receptor; *Ibabp*, ileal bile acid-binding protein; *Shp*, small heterodimer partner; TBA, total bile acids.

**Fig.S10 Effects of FGF15 on fibrosis in mouse livers.** (**A**). Liver sections were subjected to Masson's trichrome staining. Percentage of positive area was quantified with ImageJ (n=3-5). (**B**). Hepatic levels of hydroxyproline (n=3-5). (**C**). Protein level of α-SMA in the liver was analyzed by immunoblotting assay and quantified by densitometry (n=3-5). Data were shown as means ± S.E. *$p<0.05$, **$p<0.01$. α-SMA, α-smooth muscle actin.

**Fig.S11 Effects of Blu-9931 on mouse livers.** measurement of liver weight and serum ALT, AST, and ALP levels in *Atg5*-deficient livers with and without Blu-9931 (BLU) treatment. Results showed a more significant effect of BLU treatment in male mice. Data were shown as means ± S.E. *$p$<0.05, **$p$<0.01. N.S. indicates no statistical significance.  ALT, alanine transaminase;  ALP, alkaline phosphatase; AST, aspartate transaminase.

### Fig. S1

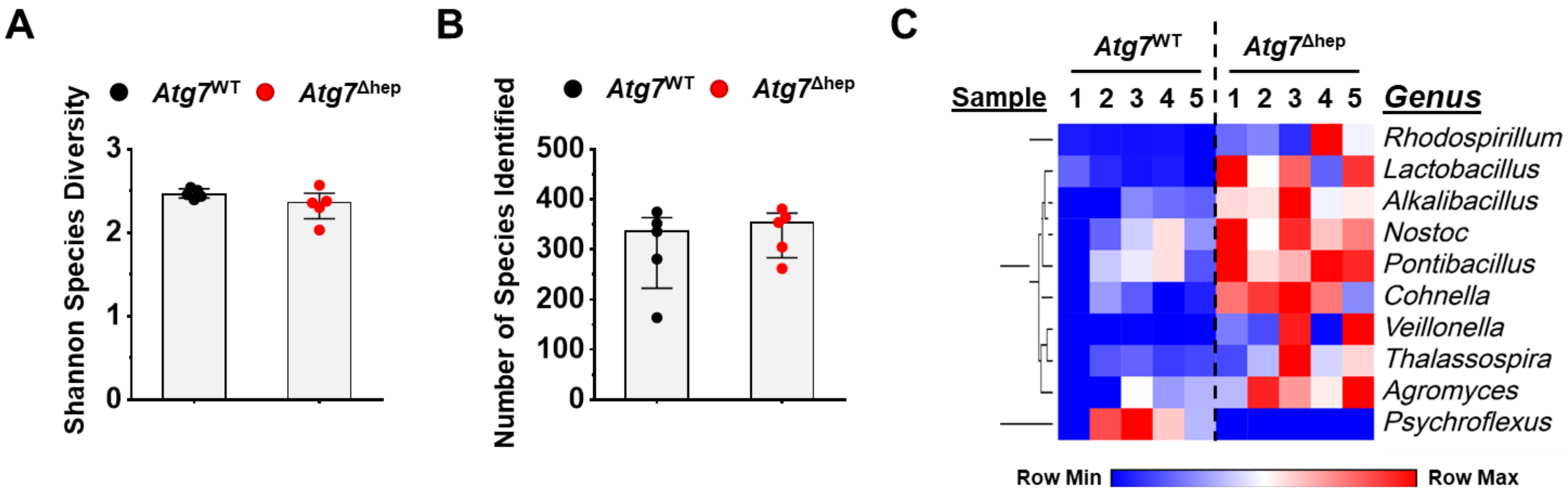

A
Shannon Species Diversity
Atg7^WT
Atg7^Δhep
3
2
1
0
B
Number of Species Identified
Atg7^WT
Atg7^Δhep
500
400
300
200
100
0
C
Atg7^WT
Atg7^Δhep
Sample
1 2 3 4 5
1 2 3 4 5
Genus
Rhodospirillum
Lactobacillus
Alkalibacillus
Nostoc
Pontibacillus
Cohnella
Veillonella
Thalassospira
Agromyces
Psychroflexus
Row Min
Row Max

#### Fig. S2

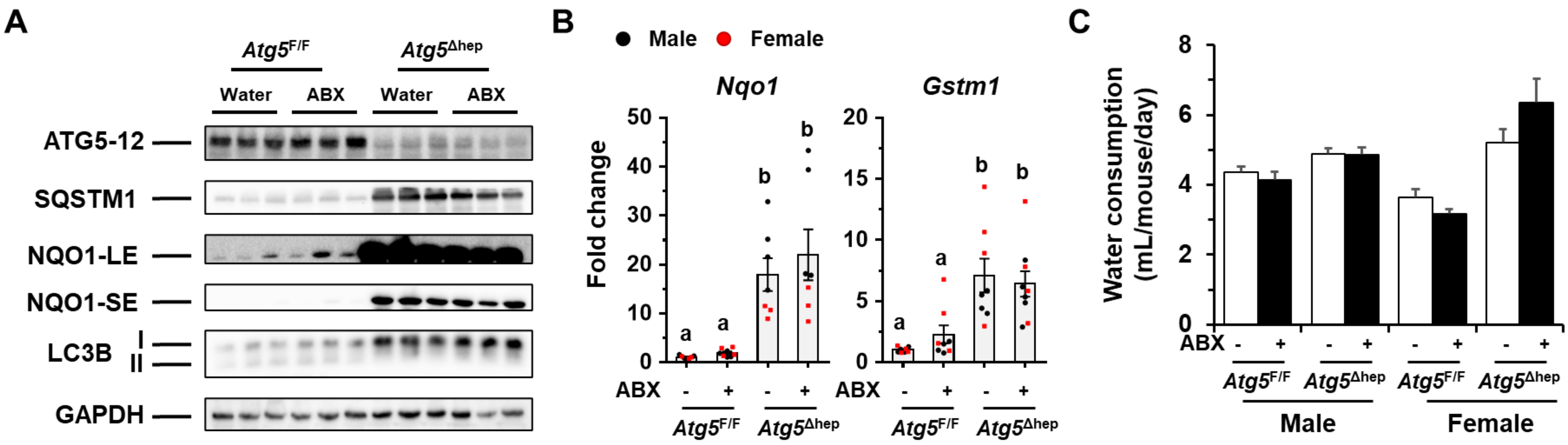
A
Atg5F/F
Atg5Δhep
Water
ABX
Water
ABX
ATG5-12
SQSTM1
NQO1-LE
NQO1-SE
LC3B
I
II
GAPDH
B
Male
Female
Nqo1
Gstm1
Fold change
ABX
Atg5F/F
Atg5Δhep
C
Water consumption (mL/mouse/day)
ABX
Atg5F/F
Atg5Δhep
Male
Female

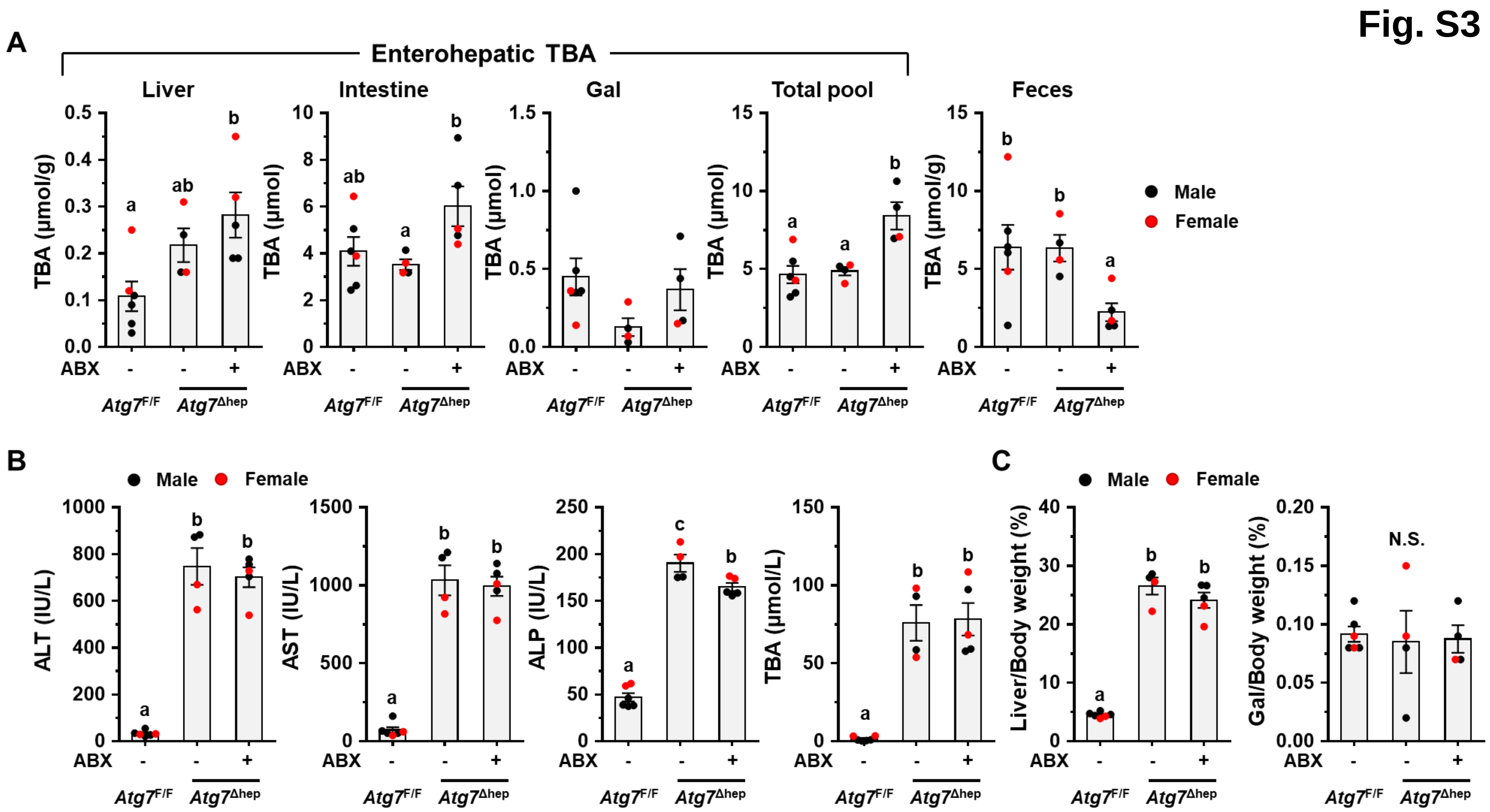
Fig. S3
A
Enterohepatic TBA
Liver
Intestine
Gal
Total pool
Feces
TBA (μmol/g)
TBA (μmol)
Male
Female
ABX
Atg7F/F
Atg7Δhep
B
ALT (IU/L)
AST (IU/L)
ALP (IU/L)
TBA (μmol/L)
C
Liver/Body weight (%)
Gal/Body weight (%)
N.S.

Fig. S4

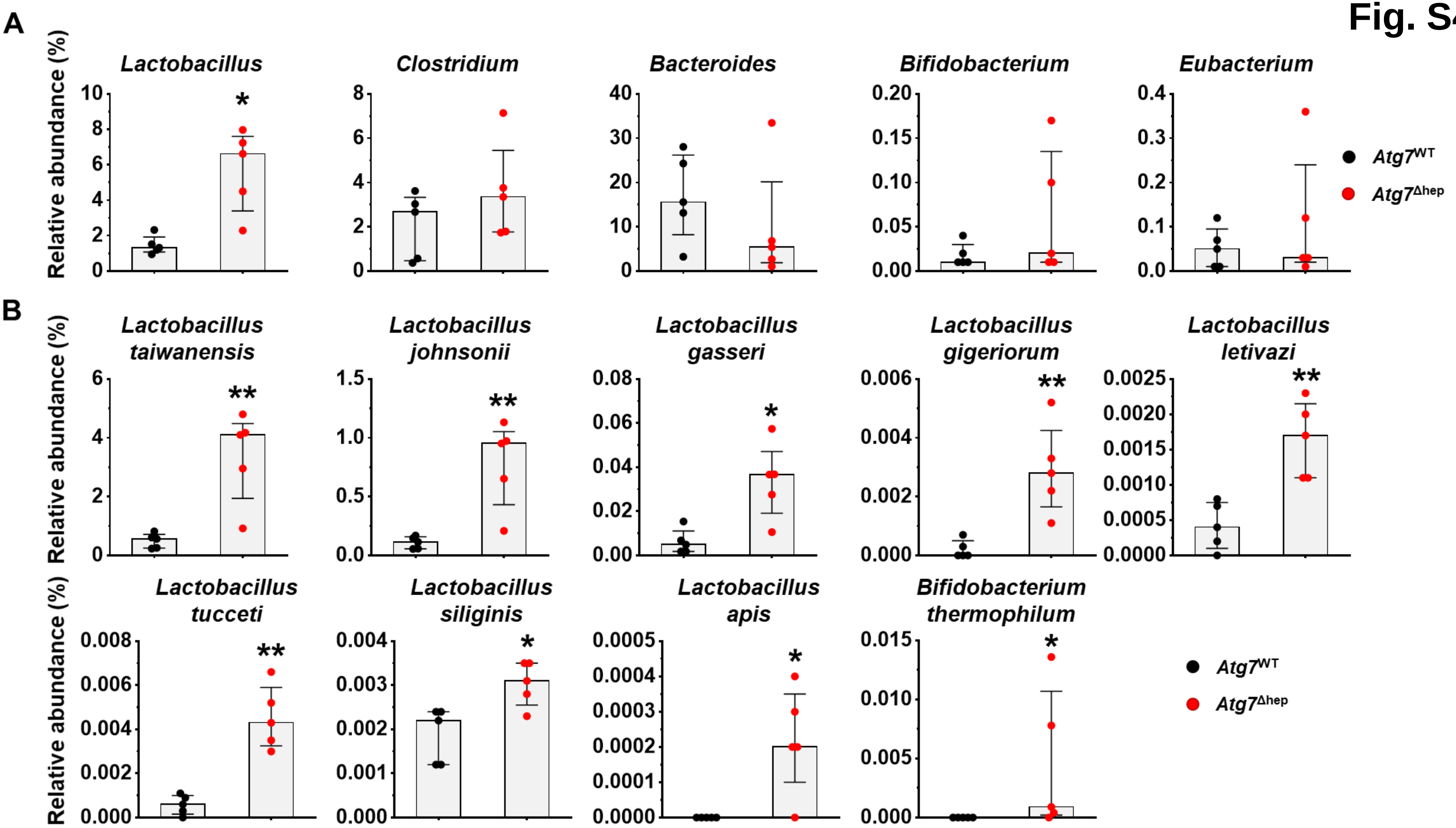
A
B
Relative abundance (%)
Lactobacillus
Clostridium
Bacteroides
Bifidobacterium
Eubacterium
Lactobacillus taiwanensis
Lactobacillus johnsonii
Lactobacillus gasseri
Lactobacillus gigeriorum
Lactobacillus letivazi
Lactobacillus tucceti
Lactobacillus siliginis
Lactobacillus apis
Bifidobacterium thermophilum
Atg7^WT
Atg7^Δhep
*
**

### Fig. S5

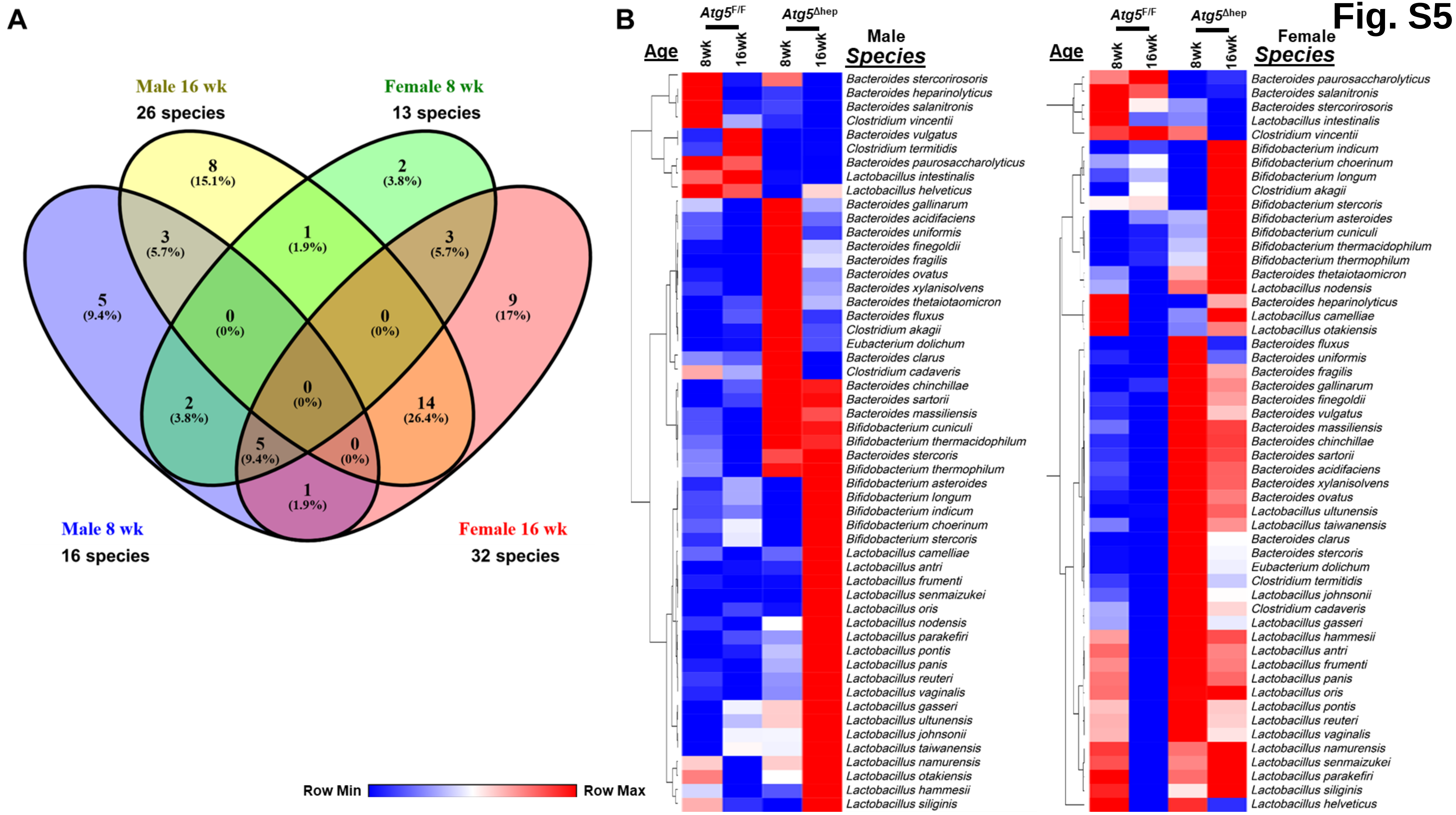
A
Male 16 wk
26 species
Female 8 wk
13 species
Male 8 wk
16 species
Female 16 wk
32 species
8 (15.1%)
2 (3.8%)
3 (5.7%)
1 (1.9%)
3 (5.7%)
5 (9.4%)
0 (0%)
0 (0%)
9 (17%)
0 (0%)
2 (3.8%)
14 (26.4%)
5 (9.4%)
0 (0%)
1 (1.9%)
B
Age
Atg5F/F
Atg5Δhep
8wk
16wk
Male
Species
Bacteroides stercorirosoris
Bacteroides heparinolyticus
Bacteroides salanitronis
Clostridium vincentii
Bacteroides vulgatus
Clostridium termitidis
Bacteroides paurosaccharolyticus
Lactobacillus intestinalis
Lactobacillus helveticus
Bacteroides gallinarum
Bacteroides acidifaciens
Bacteroides uniformis
Bacteroides finegoldii
Bacteroides fragilis
Bacteroides ovatus
Bacteroides xylanisolvens
Bacteroides thetaiotaomicron
Bacteroides fluxus
Clostridium akagii
Eubacterium dolichum
Bacteroides clarus
Clostridium cadaveris
Bacteroides chinchillae
Bacteroides sartorii
Bacteroides massiliensis
Bifidobacterium cuniculi
Bifidobacterium thermacidophilum
Bacteroides stercoris
Bifidobacterium thermophilum
Bifidobacterium asteroides
Bifidobacterium longum
Bifidobacterium indicum
Bifidobacterium choerinum
Bifidobacterium stercoris
Lactobacillus camelliae
Lactobacillus antri
Lactobacillus frumenti
Lactobacillus senmaizukei
Lactobacillus oris
Lactobacillus nodensis
Lactobacillus parakefiri
Lactobacillus pontis
Lactobacillus panis
Lactobacillus reuteri
Lactobacillus vaginalis
Lactobacillus gasseri
Lactobacillus ultunensis
Lactobacillus johnsonii
Lactobacillus taiwanensis
Lactobacillus namurensis
Lactobacillus otakiensis
Lactobacillus hammesii
Lactobacillus siliginis
Age
Atg5F/F
Atg5Δhep
8wk
16wk
Female
Species
Bacteroides paurosaccharolyticus
Bacteroides salanitronis
Bacteroides stercorirosoris
Lactobacillus intestinalis
Clostridium vincentii
Bifidobacterium indicum
Bifidobacterium choerinum
Bifidobacterium longum
Clostridium akagii
Bifidobacterium stercoris
Bifidobacterium asteroides
Bifidobacterium cuniculi
Bifidobacterium thermacidophilum
Bifidobacterium thermophilum
Bacteroides thetaiotaomicron
Lactobacillus nodensis
Bacteroides heparinolyticus
Lactobacillus camelliae
Lactobacillus otakiensis
Bacteroides fluxus
Bacteroides uniformis
Bacteroides fragilis
Bacteroides gallinarum
Bacteroides finegoldii
Bacteroides vulgatus
Bacteroides massiliensis
Bacteroides chinchillae
Bacteroides sartorii
Bacteroides acidifaciens
Bacteroides xylanisolvens
Bacteroides ovatus
Lactobacillus ultunensis
Lactobacillus taiwanensis
Bacteroides clarus
Bacteroides stercoris
Eubacterium dolichum
Clostridium termitidis
Lactobacillus johnsonii
Clostridium cadaveris
Lactobacillus gasseri
Lactobacillus hammesii
Lactobacillus antri
Lactobacillus frumenti
Lactobacillus panis
Lactobacillus oris
Lactobacillus pontis
Lactobacillus reuteri
Lactobacillus vaginalis
Lactobacillus namurensis
Lactobacillus senmaizukei
Lactobacillus parakefiri
Lactobacillus siliginis
Lactobacillus helveticus
Row Min
Row Max

**Fig. S6**

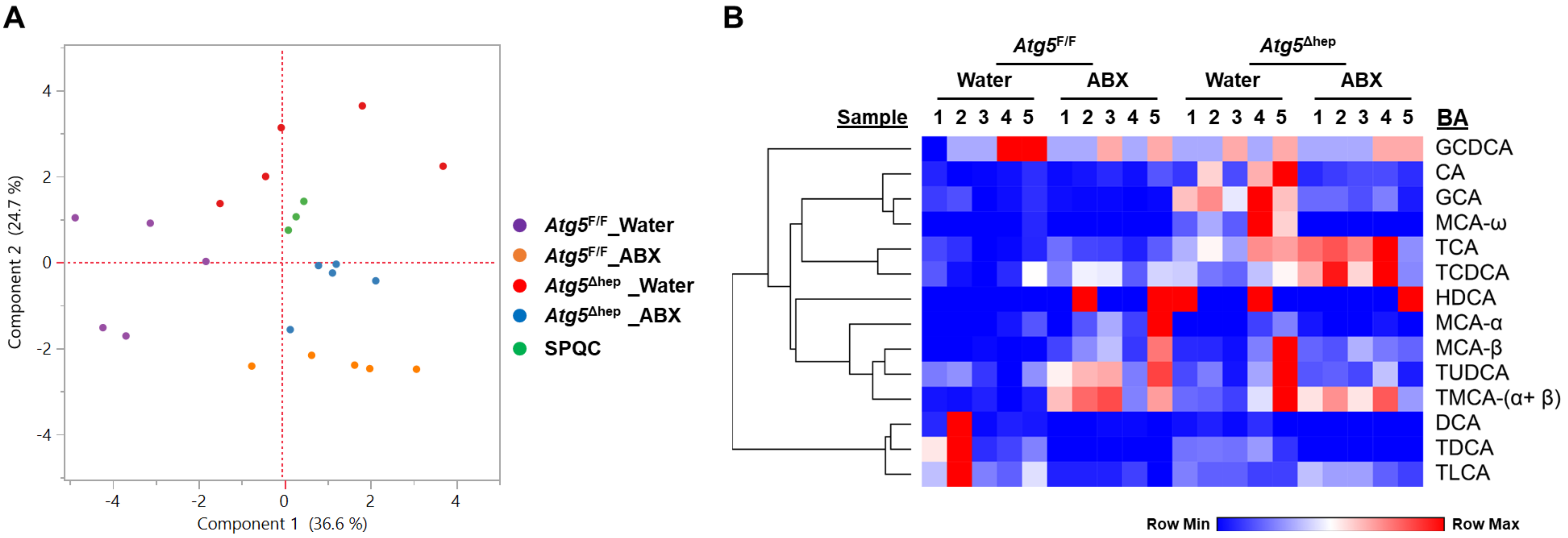
A
Component 2 (24.7 %)
Component 1 (36.6 %)
Atg5F/F_Water
Atg5F/F_ABX
Atg5Δhep _Water
Atg5Δhep _ABX
SPQC
B
Atg5F/F
Atg5Δhep
Water
ABX
Sample
1 2 3 4 5
BA
GCDCA
CA
GCA
MCA-ω
TCA
TCDCA
HDCA
MCA-α
MCA-β
TUDCA
TMCA-(α+ β)
DCA
TDCA
TLCA
Row Min
Row Max

Fig. S6

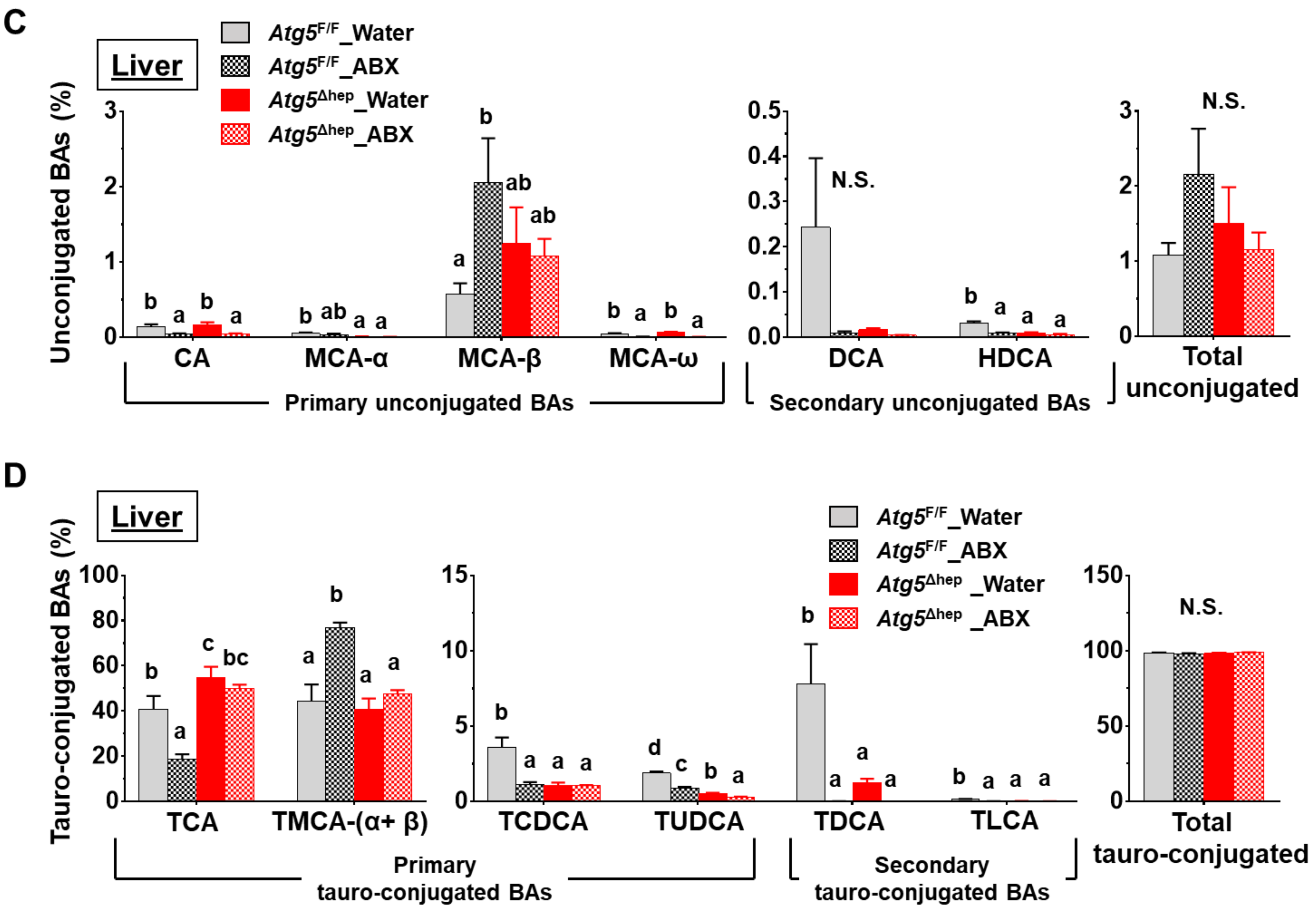
C
Liver
Atg5F/F_Water
Atg5F/F_ABX
Atg5Δhep_Water
Atg5Δhep_ABX
Unconjugated BAs (%)
CA
MCA-α
MCA-β
MCA-ω
Primary unconjugated BAs
DCA
HDCA
Secondary unconjugated BAs
N.S.
Total unconjugated
D
Liver
Tauro-conjugated BAs (%)
TCA
TMCA-(α+ β)
TCDCA
TUDCA
Primary tauro-conjugated BAs
TDCA
TLCA
Secondary tauro-conjugated BAs
Total tauro-conjugated

Fig. S7

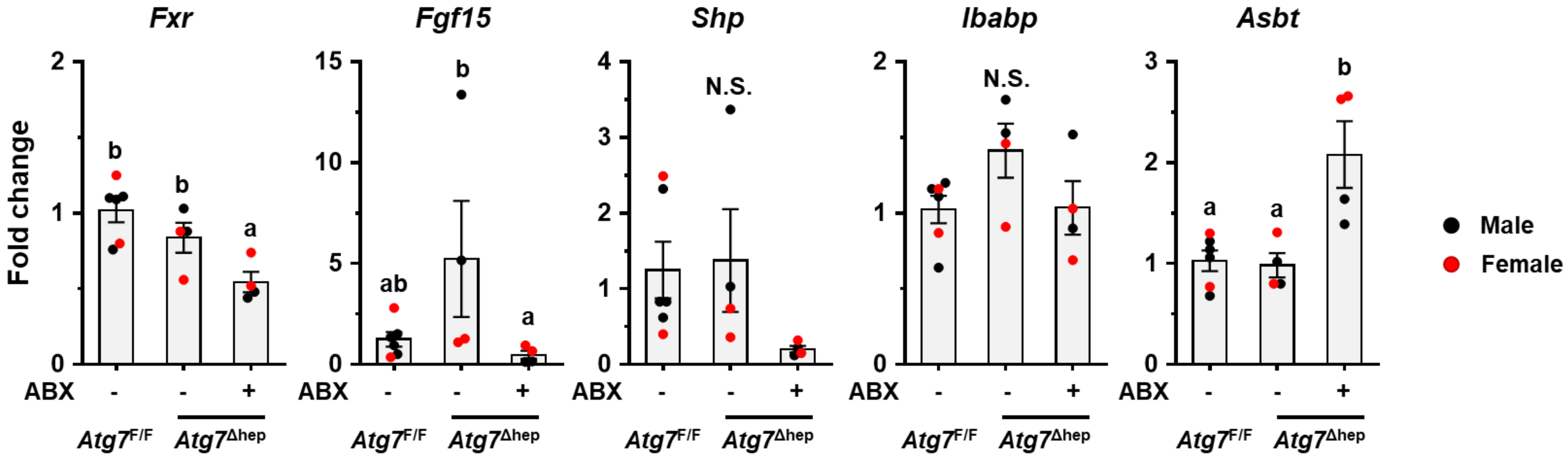

Fxr
Fgf15
Shp
Ibabp
Asbt
Fold change
b
b
a
ab
b
a
N.S.
N.S.
a
a
b
ABX
-
-
+
Atg7F/F
Atg7Δhep
Male
Female

Fig. S8

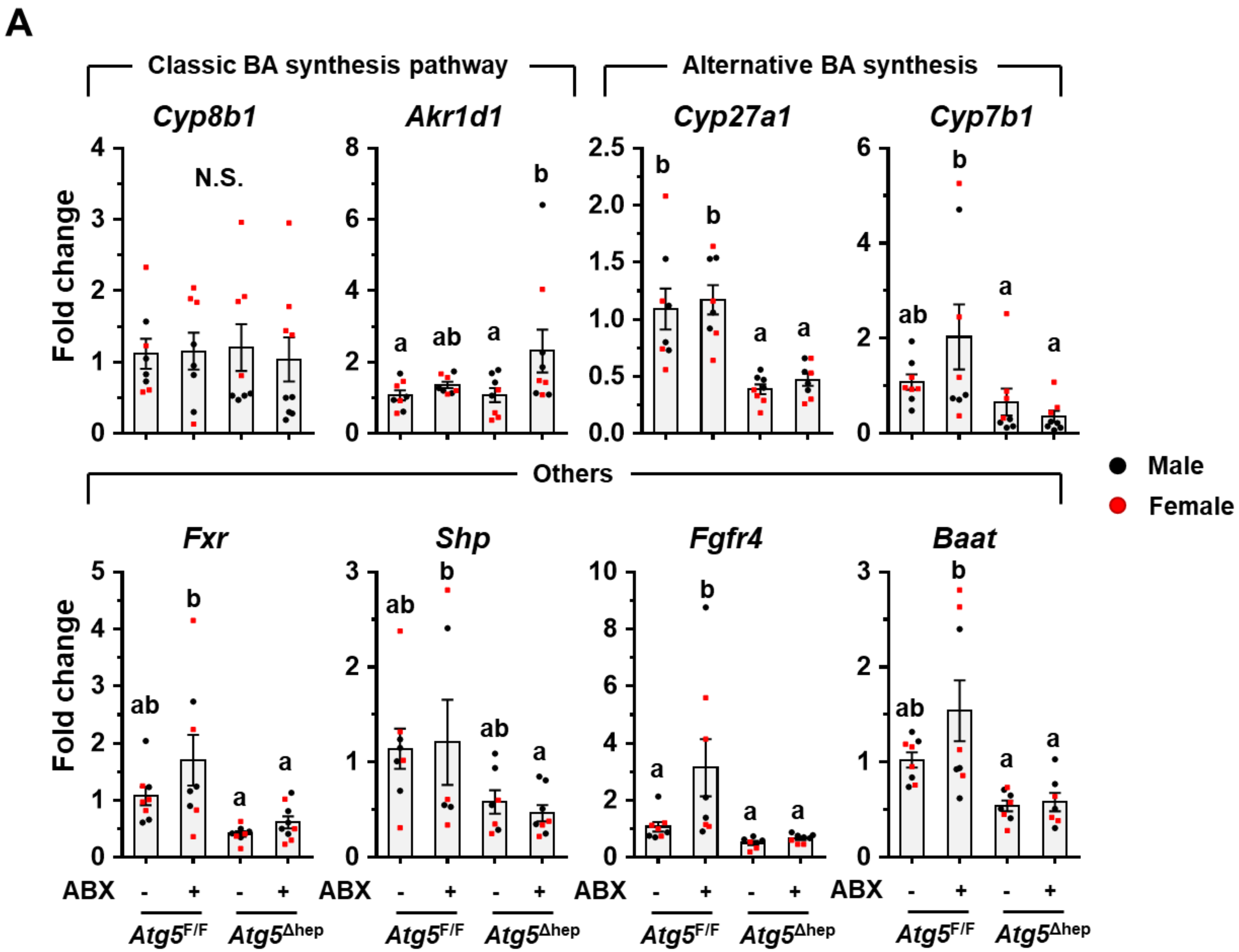

A
Classic BA synthesis pathway
Alternative BA synthesis
Cyp8b1
Akr1d1
Cyp27a1
Cyp7b1
N.S.
Fold change
Others
Male
Female
Fxr
Shp
Fgfr4
Baat
ABX
- + - +
Atg5F/F
Atg5Δhep

Fig. S8

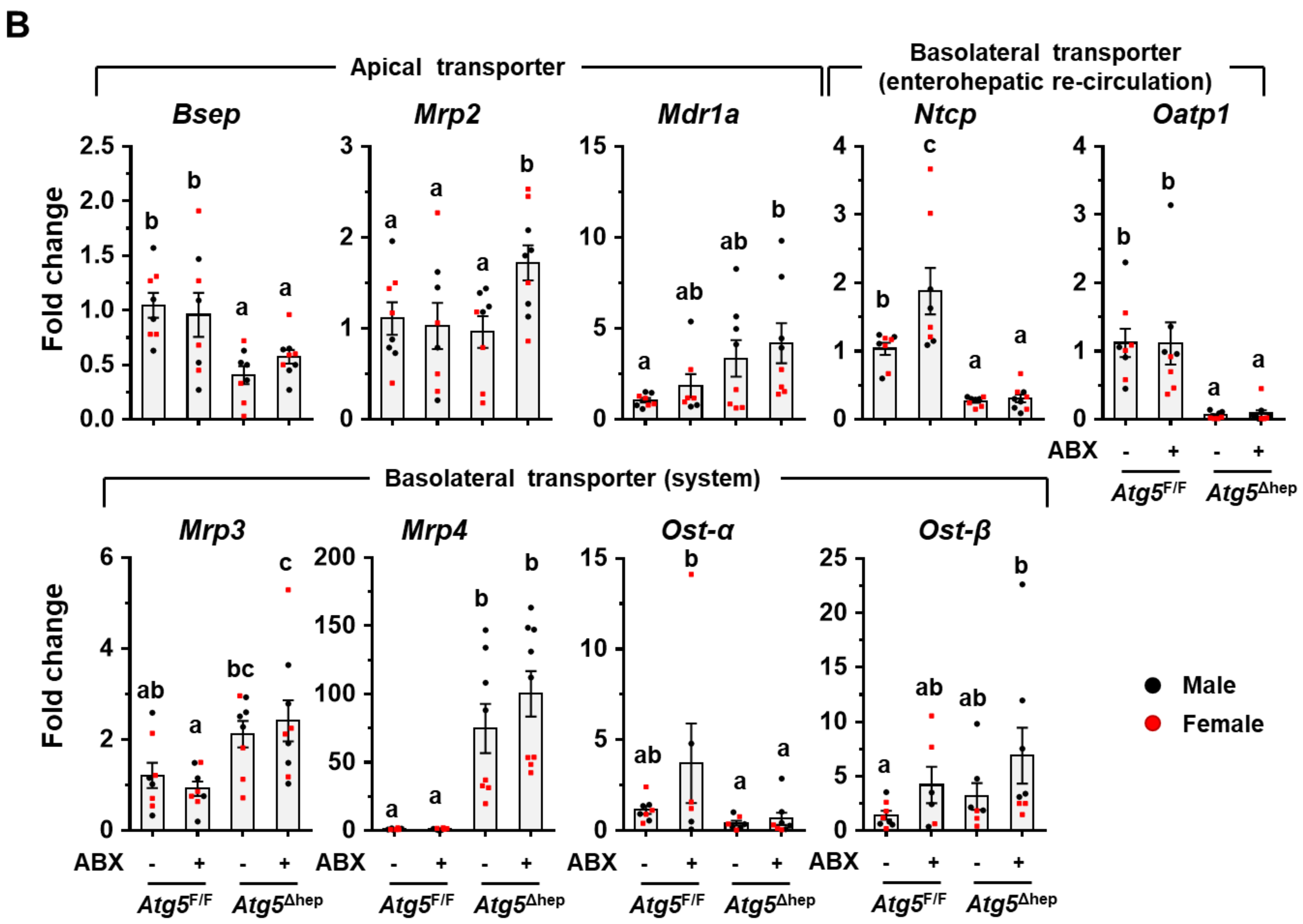
B
Apical transporter
Basolateral transporter (enterohepatic re-circulation)
Bsep
Mrp2
Mdr1a
Ntcp
Oatp1
Fold change
Basolateral transporter (system)
Mrp3
Mrp4
Ost-α
Ost-β
ABX
Atg5F/F
Atg5Δhep
Male
Female

Fig. S9

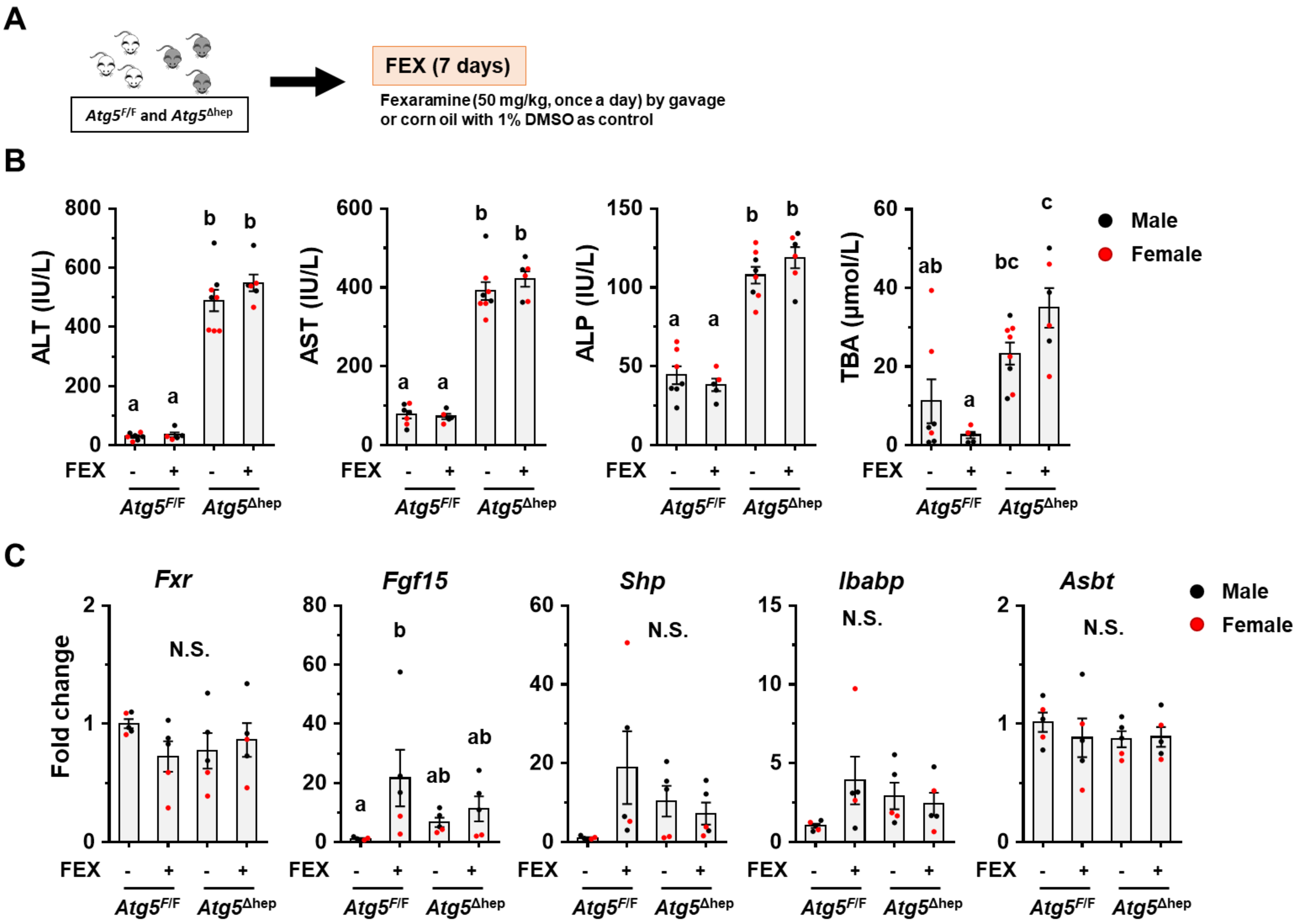
A
Atg5F/F and Atg5Δhep
FEX (7 days)
Fexaramine (50 mg/kg, once a day) by gavage
or corn oil with 1% DMSO as control
B
ALT (IU/L)
AST (IU/L)
ALP (IU/L)
TBA (μmol/L)
Male
Female
FEX
Atg5F/F
Atg5Δhep
C
Fxr
Fgf15
Shp
Ibabp
Asbt
Fold change
N.S.

Fig. S9

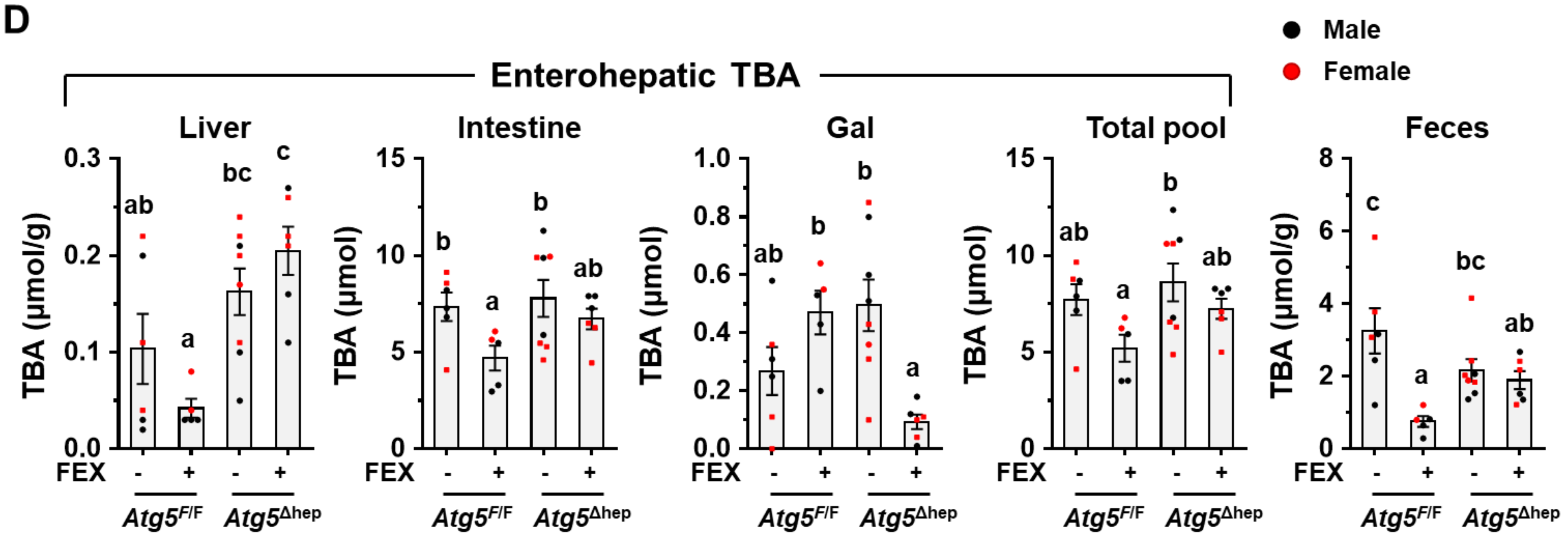
D
Male
Female
Enterohepatic TBA
Liver
Intestine
Gal
Total pool
Feces
TBA (μmol/g)
TBA (μmol)
TBA (μmol)
TBA (μmol)
TBA (μmol/g)
FEX
Atg5F/F
Atg5Δhep

Fig. S10

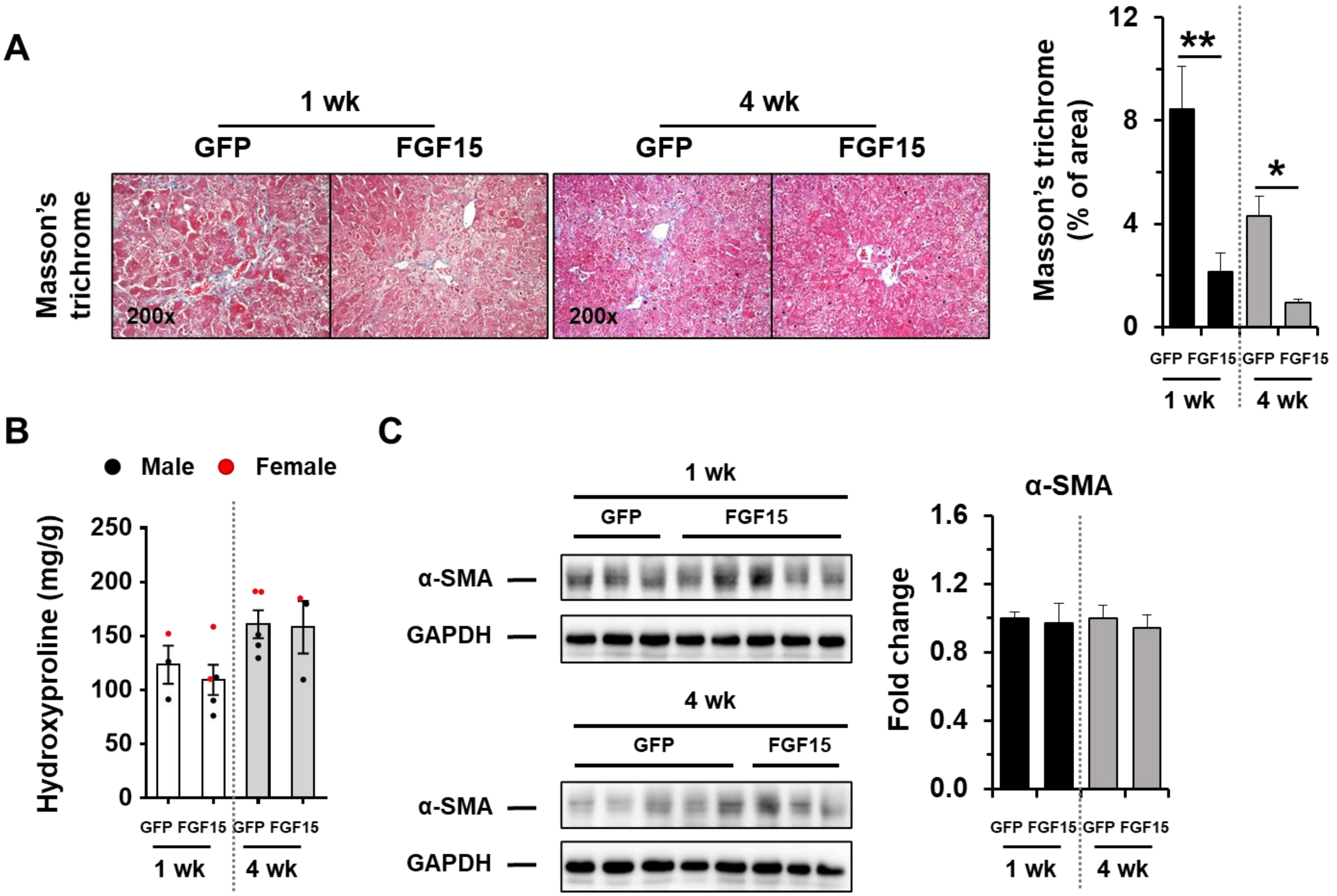

A
1 wk
4 wk
GFP
FGF15
GFP
FGF15
Masson's trichrome
200x
200x
Masson's trichrome (% of area)
12
8
4
0
**
*
GFP FGF15 GFP FGF15
1 wk
4 wk
B
Male
Female
Hydroxyproline (mg/g)
250
200
150
100
50
0
GFP FGF15 GFP FGF15
1 wk
4 wk
C
1 wk
GFP
FGF15
α-SMA
GAPDH
4 wk
GFP
FGF15
α-SMA
GAPDH
α-SMA
Fold change
1.6
1.2
0.8
0.4
0.0
GFP FGF15 GFP FGF15
1 wk
4 wk

**Fig. S11**

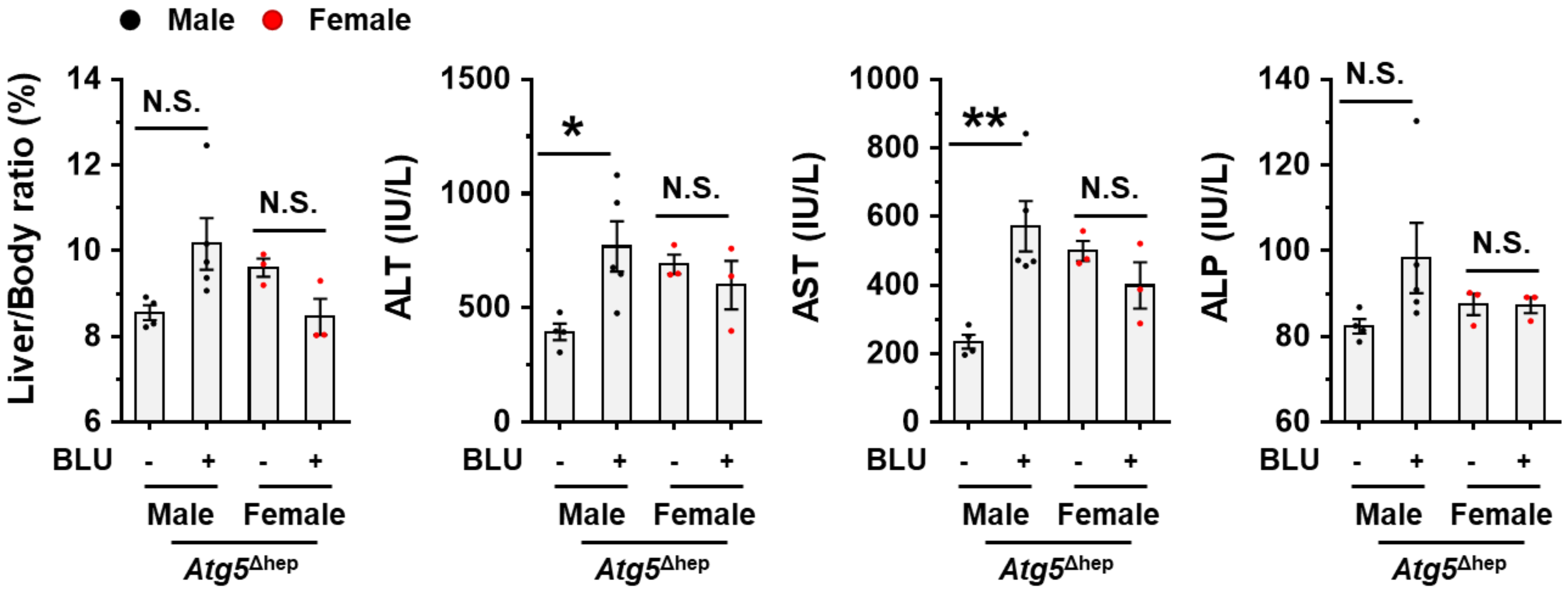
Male
Female
Liver/Body ratio (%)
14
12
10
8
6
N.S.
N.S.
BLU
-
+
-
+
Male
Female
Atg5^Δhep
ALT (IU/L)
1500
1000
500
0
*
N.S.
AST (IU/L)
1000
800
600
400
200
0
**
N.S.
ALP (IU/L)
140
120
100
80
60
N.S.
N.S.
